# Dynamic brain states during encoding and their post-encoding reinstatement predict episodic memory in children

**DOI:** 10.1101/2025.11.25.690570

**Authors:** Yimeng Zeng, Sandhya P. Chakravartti, Srikanth Ryali, Lei Li, Shaozheng Qin, Vinod Menon

## Abstract

Episodic memory, the ability to rapidly learn and explicitly remember past events and experiences, plays a critical role in children’s academic learning and knowledge acquisition. The formation of lasting memories relies on the brain’s ability to dynamically organize its activity. How these neural configurations unfold moment-by-moment across encoding and offline phases remains poorly understood. To probe these dynamics, we applied a novel Bayesian Switching Dynamic Systems approach, a hidden Markov model with automatic state detection, to fMRI data from children performing scene encoding followed by an offline post-encoding rest. We identified four distinct brain states during encoding with unique activation modes between visual, medial temporal lobe, and frontoparietal and default mode network nodes. An “active-encoding” state with integrated visual-hippocampal and frontoparietal activity dominated encoding and predicted individual memory performance, while an inactive state negatively predicted performance. State transition dynamics revealed that flexible shifts into the active-encoding state enhanced memory formation, whereas transitions toward inactive states impaired it, demonstrating that memory success depends on dynamic neural flexibility. Critically, encoding states spontaneously reemerged during post-encoding rest. A “default-mode” state characterized by enhanced default mode network activity showed sustained maintenance during rest and robustly predicted memory outcomes, an effect specific to post-encoding, not pre-encoding rest. These findings establish that episodic memory emerges from coordinated brain state sequences bridging online encoding with offline consolidation, providing a computational framework for how moment-to-moment neural dynamics support memory formation in children.

**Significance Statement:** How do children’s brains form lasting memories? Using a novel computational approach, we reveal that memory formation depends on the brain’s ability to rapidly shift between distinct coordination patterns, or “brain states.” During learning, children who flexibly transition into an optimal brain state form stronger memory. Remarkably, these encoding states spontaneously reactivate during subsequent rest, and children who sustain specific default-mode network states longer show better memory retention. These findings demonstrate that successful memory emerges from dynamic neural flexibility across both active encoding and offline rest, offering a new computational framework for understanding memory formation in the developing brain.

## Introduction

Understanding how children’s brains form lasting memories is a fundamental question in developmental cognitive neuroscience ^1–3^. Episodic memory, the ability to rapidly learn and explicitly remember past events and experiences ^4^, plays a critical role in children’s academic learning and knowledge acquisition ^2,5^. Episodic memory formation requires the brain to coordinate activity across multiple distributed systems through flexible, moment-to-moment functional interactions during both online encoding and subsequent offline consolidation phases ^6–9^. Despite its developmental importance, the dynamic neural mechanisms underlying this essential cognitive capacity remain unclear. Here, using a novel computational approach that identifies transient coordination patterns, or “brain states”, we examine how the developing brain dynamically shifts between these states during learning and how these states are spontaneously reinstated during rest to support memory consolidation. Characterizing these dynamic processes is critical for identifying the neural mechanisms that support learning and memory during development.

Decades of research have identified key brain regions essential for episodic memory formation, revealing a hierarchical processing architecture ^10,11^. Visual processing begins in ventral stream pathways from the occipital lobe, which extract visual features and object representations from incoming stimuli. This perceptual information is transmitted to the medial temporal lobe (MTL), particularly the hippocampus and parahippocampal gyrus, which serves as the critical binding hub for integrating diverse perceptual, spatial, and temporal elements into coherent, detailed episodic representations ^12–15^, especially vital for children’s developing episodic memory abilities^1^. Beyond these sensory and medial temporal systems, the lateral frontoparietal network (FPN) provides top-down cognitive control and attentional modulation that selectively guides which information enters memory and how encoding processes unfold across time ^16–18^. This network undergoes substantial developmental changes, making it a particularly important contributor to episodic memory encoding and consolidation during childhood^19^. The default mode network (DMN), particularly its medial prefrontal cortex (mPFC) and posterior cingulate cortex (PCC) nodes, also plays a critical role in episodic memory formation. DMN regions are thought to undergo flexible reconfiguration, transiently integrating with task-positive networks during encoding to support self-referential processing and contextual integration^20,21^ and may function as a coordinated network to support episodic memory development ^22^. However, the precise neural dynamics by which these distributed brain systems interact to support episodic memory formation in children remain poorly understood.

While prior research has provided important insights into how episodic memory relies on individual brain regions, successful episodic memory formation critically depends on the temporal coordination between these systems. Brain regions must coordinate their activity through dynamic functional interactions, rapidly reconfiguring their activity and connectivity patterns in response to changing cognitive demands and stimulus characteristics. This dynamic network reconfiguration (dynamic state) enables flexible adaptation to diverse task contexts – a capacity that is fundamental for efficient learning ^23–25^. Investigations of memory systems in children have traditionally focused on the role of individual brain regions ^1–3^, but how these systems interact dynamically during encoding and how they coordinate during offline consolidation to support memory formation remains largely unknown.

Fundamental questions remain unanswered: how do distributed brain regions dynamically assemble into specific functional states during encoding? Whether the frequency of state emergence (occupancy rate) or its temporal duration (lifetime) could be associated with episodic memory encoding? How do transitions between brain states determine memory success? Understanding these dynamic assembly processes is critical because episodic memory formation likely depends on the brain’s ability to flexibly coordinate distributed circuits into optimal network configurations to support memory formation, rather than sustained activation of individual regions. Conventional functional connectivity analyses and regional activation studies cannot detect the dynamic coordination underlying episodic memory, creating critical knowledge gaps. Novel computational approaches are needed to identify brain states that reflect the dynamic assembly of distributed neural circuits.

A second critical gap concerns the relationship between encoding dynamics and offline consolidation processes. Recent research has demonstrated that during awake rest or sleep, specific neural circuits automatically retrieve and reinstate memory traces – a process known as replay or reinstatement that is crucial for memory consolidation ^6,7,26^. This reinstatement likely requires extensive dynamic coordination between brain areas to facilitate reorganization of memory representations into long-term storage ^7,27,28^. However, a fundamental question remains: Do the specific brain states that emerge during encoding spontaneously re-occur during post-encoding rest, and does this reinstatement predict memory outcomes? Understanding how dynamic interactions among memory-related brain states evolve from encoding to post-encoding rest is critical for revealing the neural mechanisms underlying memory formation. This represents a key missing piece in our understanding of how episodic memories are formed, maintained, and consolidated across online and offline phases. Previous studies in adults have reported that persistent brain activity could occur during post-encoding rest and core brain regions’ communications of hippocampal–mPFC coherence could be critical for episodic memory consolidation ^29,30^. Specific neural processes such as replay and reactivation may serve as key mechanisms for offline episodic memory consolidation ^6,7^. However, whether similar phenomenon occur in children still remains largely unknown.

To address these critical gaps, we implemented a novel Bayesian Switching Dynamic Systems (BSDS) approach combined with graph-theoretic network analyses. BSDS implements a hidden Markov model with automatic relevance detection that determines the optimal number of brain states through unsupervised learning, overcoming the arbitrary parameter selection that constrains conventional approaches ^31^. The BSDS model offers several key advantages: it identifies latent brain states and their temporal evolution without arbitrary parameter constraints, quantifies state lifetime and occurrence rates, and assesses dynamic transition patterns using moment-to-moment detection algorithms. Crucially, BSDS can probe mental processes that are internally organized rather than strictly constrained by external stimuli, making it ideally suited for capturing flexible, adaptive neural dynamics underlying memory formation ^31,32^.

Twenty-four typically developing children (ages 8-13) underwent an fMRI scan while performing episodic memory encoding of natural scenes, followed by a 6-minute post-encoding rest session and a subsequent recognition memory test with confidence ratings outside the scanner (**Figure 1**). We focused on 16 core regions spanning visual cortex, medial temporal lobe, frontoparietal networks, and DMN nodes that showed significant subsequent memory effects (remembered > forgotten contrast). We tested three primary hypotheses. First, BSDS would reveal dissociable brain states with distinct functional connectivity patterns among memory-related circuits during encoding, with occurrence of specific encoding-optimal states predicting individual memory performance. Second, dynamic transitions between brain states would be critical for memory formation, with transitions toward optimal encoding states enhancing performance and transitions toward suboptimal states impairing it. Third, and most importantly, brain states identified during encoding would spontaneously re-occur during post-encoding rest through reinstatement mechanisms, with offline state reactivation predicting memory consolidation success. This would provide direct evidence for systems consolidation through dynamic state reinstatement.

**Figure 1.**
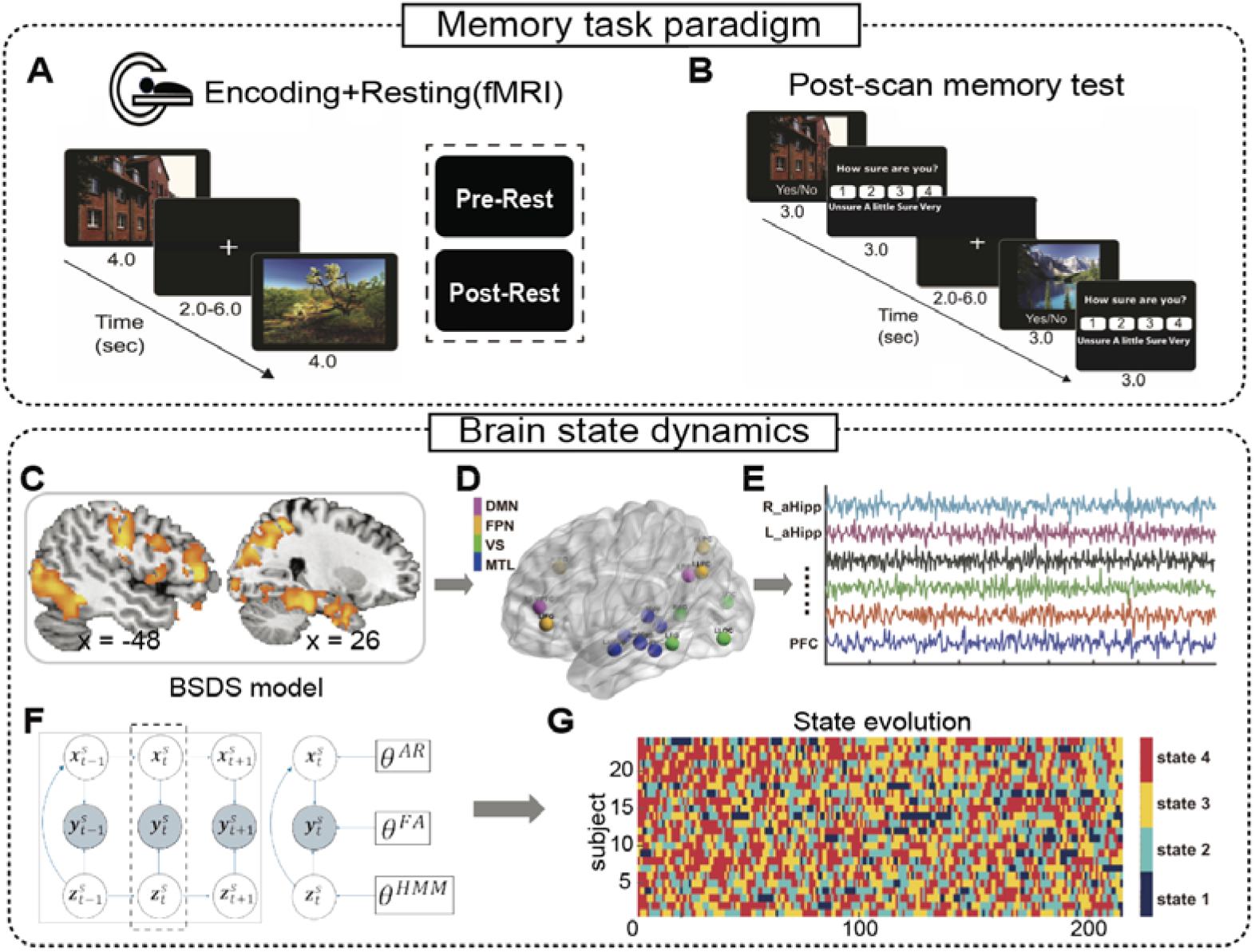
Experimental design and dynamic brain state modeling framework. **(A) episodic memory paradigm.** During fMRI scanning, participants encoded 100 images of buildings and natural scenes (3 seconds each, jittered fixation 2-6 seconds), followed by a 6-minute post-encoding rest period. Participants were instructed to imagine themselves in each scene and judge whether it was natural or building. **(B) Post-scan memory assessment.** Outside the scanner, participants completed a memory retrieval test with confidence ratings using a 4-point scale (“Unsure” to “Very Sure”). **(C) Memory-related brain regions.** Lateral view showing significant activation clusters from the subsequent memory contrast (remembered > forgotten contrast) across ventral visual, medial temporal lobe, and frontal-parietal regions. **(D) Regions of Interest for state space modeling.** Sixteen ROIs spanning visual association cortex (VS: fusiform, lateral occipital complex), medial temporal lobe (MTL: hippocampus, parahippocampal cortex), frontoparietal network (FPN), and default mode network (DMN: medial prefrontal and posterior cingulate cortex). ROIs are color-coded by system. **(E) Representative fMRI time series.** Preprocessed and normalized signals from selected ROIs (labeled: right and left anterior hippocampus, medial prefrontal cortex) during encoding. **(F) Bayesian Switching Dynamic Systems model. y^*s*^_*t*_** denotes the observed ROI time-series vector at time (t) for subject (s) that parameterized by ***θ^FA^***. ***z^s^_t_*** denotes the latent states that parameterized by ***θ^HMM^***. ***X^s^_t_*** are the latent space variables that parameterized by ***θ^AR^* (G) Brain state evolution.** Color-coded temporal sequence of four BSDS-inferered brain states across all participants during encoding (first-half TRs shown), illustrating moment-to-moment state transitions.

Our computational framework provides the first quantitative characterization of how dynamic brain states bridge online encoding with offline consolidation to support episodic memory formation in children. By revealing the specific brain state configurations and transitions that optimize learning, including the critical role of DMN-mediated consolidation states during rest, this work advances our understanding of how dynamic brain state configurations support memory function in children.

## Results

### Episodic memory performance

We first confirmed that our experimental paradigm successfully engaged episodic memory processes in children (**Methods**). During fMRI scanning, participants encoded 100 images of buildings and scenes, followed by a post-scan memory retrieval test with confidence ratings (**Fig. 1A-1B**). We analyzed hit rates, confidence ratings, and their relationship to assess episodic memory formation. Children demonstrated successful episodic memory encoding with a significant hit rate (t_(23)_ = 3.97, *p* < 0.001) and they rated remembered items with higher confidence rating than forgotten ones (t_(23)_ = 7.50, *p* < 0.001) (**Fig. S1**). One-way analysis of variance (ANOVA) revealed a significant main effect of confidence on memory retrieval (F_(1,47)_ = 15.03, *p* < 0.001), with the strongest effect for highest confidence trials. These results confirm successful episodic memory encoding and confidence-dependent retrieval in our sample.

### Dynamic brain states during the episodic memory encoding phase

Episodic memory formation requires coordinated activity across distributed brain regions, yet conventional approaches cannot capture the dynamic functional interactions that support memory encoding. We next applied Bayesian Switching Dynamic Systems (BSDS), a validated state-space modeling approach ^19,31,32^, to time series from 16 memory-related regions of interest, identified through subsequent memory effects (remembered > forgotten contrast; **Fig. 1C-D**). These regions spanned four characteristic functional modules including: visual-related ventral temporo-occipital cortex, medial temporal lobe system, and frontal-parietal networks area as well as default mode networks ^33,34^ (**Supplementary Table S1**). BSDS identified latent brain states with distinct amplitude patterns and functional connectivity signatures during episodic memory encoding.

BSDS first inferred four latent brain states with unique characteristics (**Fig. 2A**). State 3 (“active-encoding state”) exhibited strong co-activation across visual areas, MTL system, and parietal regions while State 2 (“default-mode state”) showed peak activation in medial prefrontal cortex (MPFC) and posterior cingulate cortex (PCC). State 4 (“inactive state”) displayed uniformly low amplitude across all regions, while State 1 showed intermediate activation patterns.

**Figure 2.**
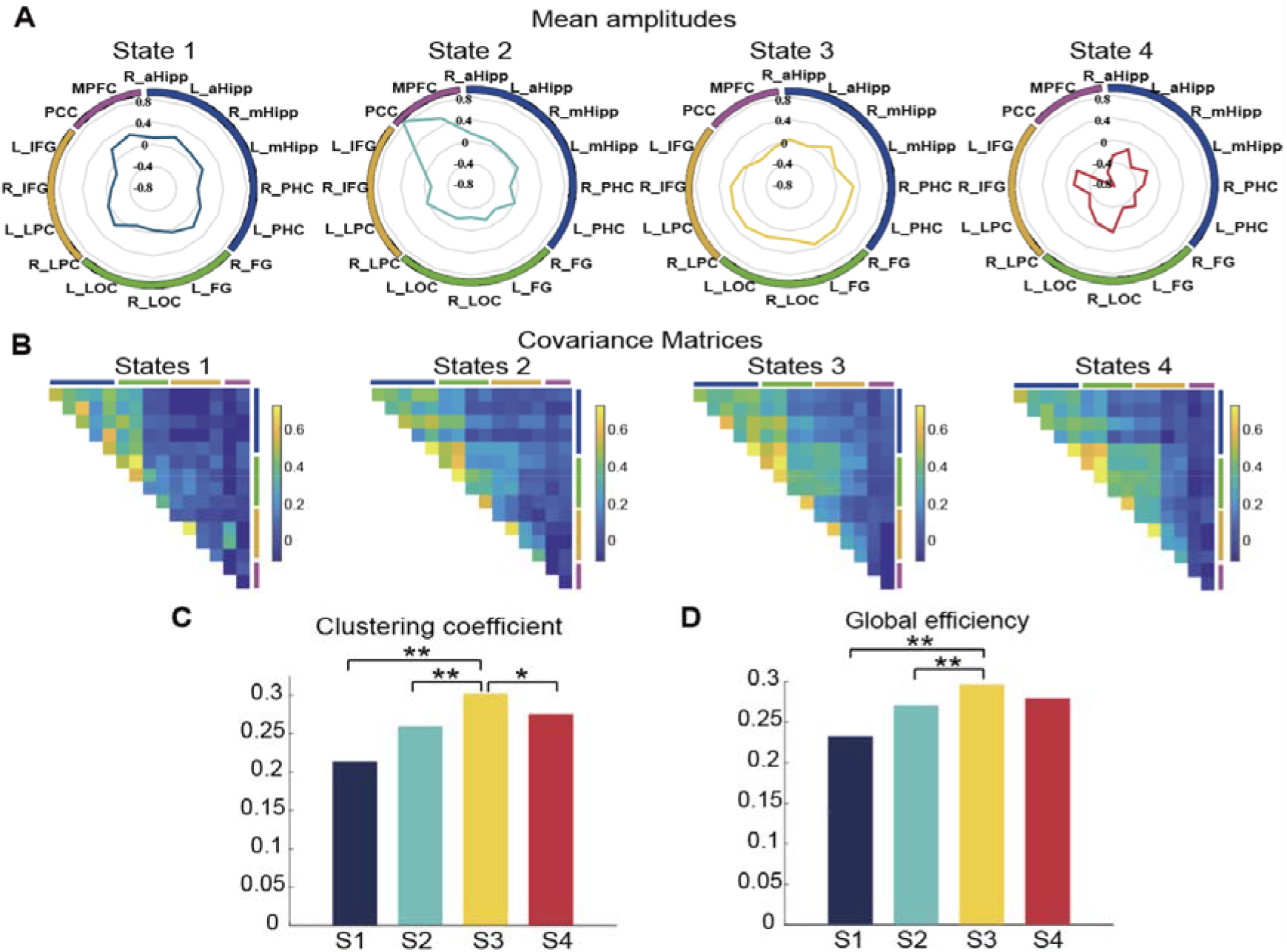
Characterization of dynamic brain states during memory encoding. **(A) State-specific activation profiles.** Polar plots depicting mean amplitude profiles across 16 ROIs for each of the four BSDS-identified brain states. State 3 (active-encoding state) shows prominent activation across visual areas, MTL system, and frontoparietal regions. State 2 (default-mode state) exhibits selective activation in mPFC and PCC. State 4 (inactive state) displays uniformly low activation, while State 1 shows intermediate activation. **(B) Functional connectivity.** Group-level covariance matrices for each brain state. The active-encoding state (S3) exhibits the highest overall functional connectivity, significantly exceeding all other states. **(C-D) Network topology.** Clustering coefficient **(C)** and global efficiency **(D)** derived from state-specific connectivity matrices. S3 shows significantly higher values on both metrics compared to all other states. Error bars represent SEM. \**p* < 0.05, \*\**p* < 0.01.

Subsequent functional connectivity analysis revealed that the active-encoding state (S3) exhibited significantly higher average connectivity during encoding (0.28) than all other states (paired *t*-tests: all *ps* < 0.001, FDR corrected; **Fig. 2B**). Graph theory analyses further demonstrated that the active-encoding state (S3) had significantly higher clustering coefficient and global efficiency compared to other states (all permuted *ps* < 0.05, FDR corrected; **Fig. 2C-D**).

These findings reveal four distinct brain state configurations during encoding, with the active-encoding state exhibiting optimal network integration and segregation properties consistent with coordinated visual processing, hippocampal binding, and attentional control.

### Brain state dynamics during the encoding phase predict memory performance

To test whether these brain states reflect behaviorally meaningful neural configurations, we next examined how their temporal dynamics, captured by state occupancy rates (OR), mean lifetime and state-to-state transition patterns, predict individual memory performance (**Methods**). We first found that identified brain states showed markedly different occupancy patterns during encoding (**Fig. 3A**). The active-encoding state (S3) dominated with 38.39 ± 7.58% occupancy rate, significantly higher than all other states (t_(23)_ > 5.43, *ps* < 0.001, FDR corrected). Meanwhile, the mean lifetimes of the four states did not differ (all ts_(23)_ < 1.47, *ps* > 0.86, FDR corrected). Critically, active-encoding state occupancy rate correlated positively with memory accuracy (r = 0.47, *p* < 0.05, FDR corrected), while inactive state occupancy showed a strong negative correlation (r = -0.64, *p* < 0.01, FDR corrected; **Fig. 3C**).

**Figure 3.**
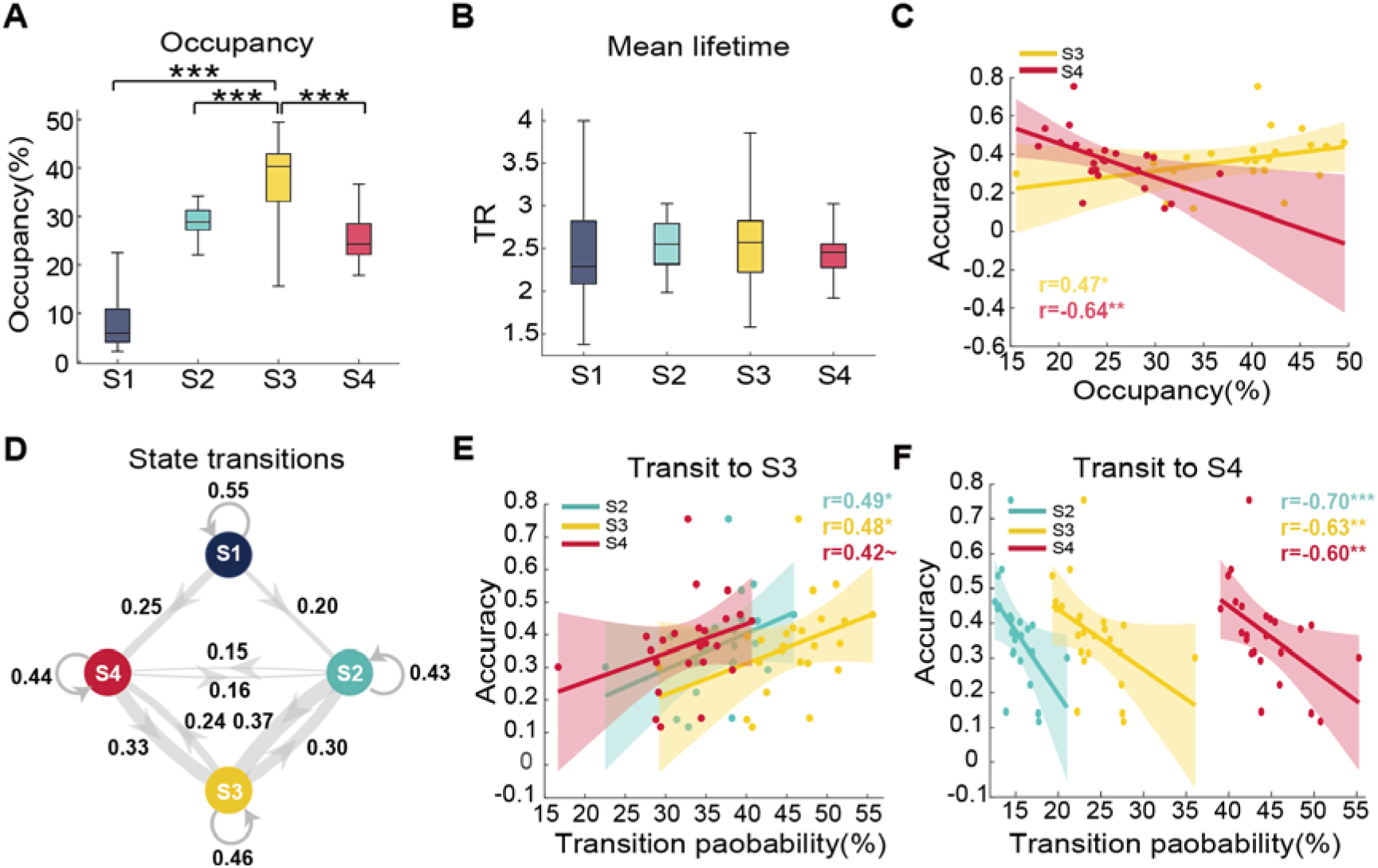
Brain state dynamics during encoding predict individual memory performance. **(A-B) State temporal properties.** Occupancy rates and mean lifetimes for each brain state during encoding. The active-encoding state (S3) dominates with 38.39% occupancy, significantly higher than all other states. **(C) State–behavior correlations.** Active-encoding state (S3) occupancy correlates positively with memory accuracy, while inactive state (S4) occupancy correlates negatively. **(D) State transition network.** Transition probabilities between brain states during encoding, with edge thickness representing transition strength. Prominent bidirectional transitions occur between active-encoding (S3), control (S2), and inactive (S4) states. **(E-F) Transition-behavior relationships.** Transitions toward the active-encoding state (S2→S3; S4→S3) enhance memory performance (**E**), while transitions toward the inactive state (S2→S4; S3→S4) impair performance (**F**). Self-transitions of S4 also negatively predict memory. Dashed lines indicate 95% confidence intervals; solid lines show best linear fit. ∼*p* < 0.10, \**p* < 0.05, \*\**p* < 0.01, \*\*\**p* < 0.001.

Further state transition analysis revealed complex dynamics with prominent transitions toward active-encoding states (**Fig. 3D**). Notably, transitions to the active-encoding state were beneficial: both default-mode state (S2) → active-encoding (r = 0.49, *p* < 0.05, FDR corrected) and inactive state (S4) → active-encoding transitions (r = 0.48, *p* < 0.1, FDR corrected) correlated positively with memory accuracy (**Fig. 3E**). Conversely, transitions toward the inactive state (S4) were detrimental, with incoming transitions showing negative correlations with memory accuracy (default-mode state (S2) → inactive (S4): r = -0.70, *p* < 0.001; active encoding → inactive: r = -0.63, *p* < 0.01, FDR corrected; **Fig. 3F**). Notably, these brain-memory associations were replicated with D-prime as alternative memory performance index (**Supplement Table. S2**).

Finally, as a control analysis, we applied BSDS to temporally permuted encoding time series, which preserved spatial co-activation patterns but disrupted event-related temporal structure. This analysis yielded no significant brain–behavior correlations (**Supplementary Figure S7),** confirming that the state dynamics captured by BSDS reflect temporally structured neural processes tied to event-related encoding rather than noise or static spatial patterns.

These results demonstrate that successful episodic memory encoding depends not only on occupying optimal brain states but also on dynamic transitions between states. The active-encoding state serves as a “hub” configuration: transitions toward it enhance memory, while transitions away from it impair encoding.

### Recurrence of latent encoding brain states during the post-encoding rest phase supports memory consolidation

Systems consolidation models propose that memory traces are reactivated during subsequent rest periods to facilitate long-term storage ^35–37^. Next, we examined whether brain states identified during encoding would re-emerge during post-encoding rest, and whether this reinstatement would be associated with memory performance. We applied the BSDS model trained on encoding data to decode brain states during the pre- and post-encoding rest session using maximum log-likelihood estimation (**Fig. 4A, Methods**). We also confirmed that the overall state decoding strength was significantly greater than values under the null distribution (permutation-*p* < 0.05) (**Supplementary Figure S3, Methods**).

**Figure 4.**
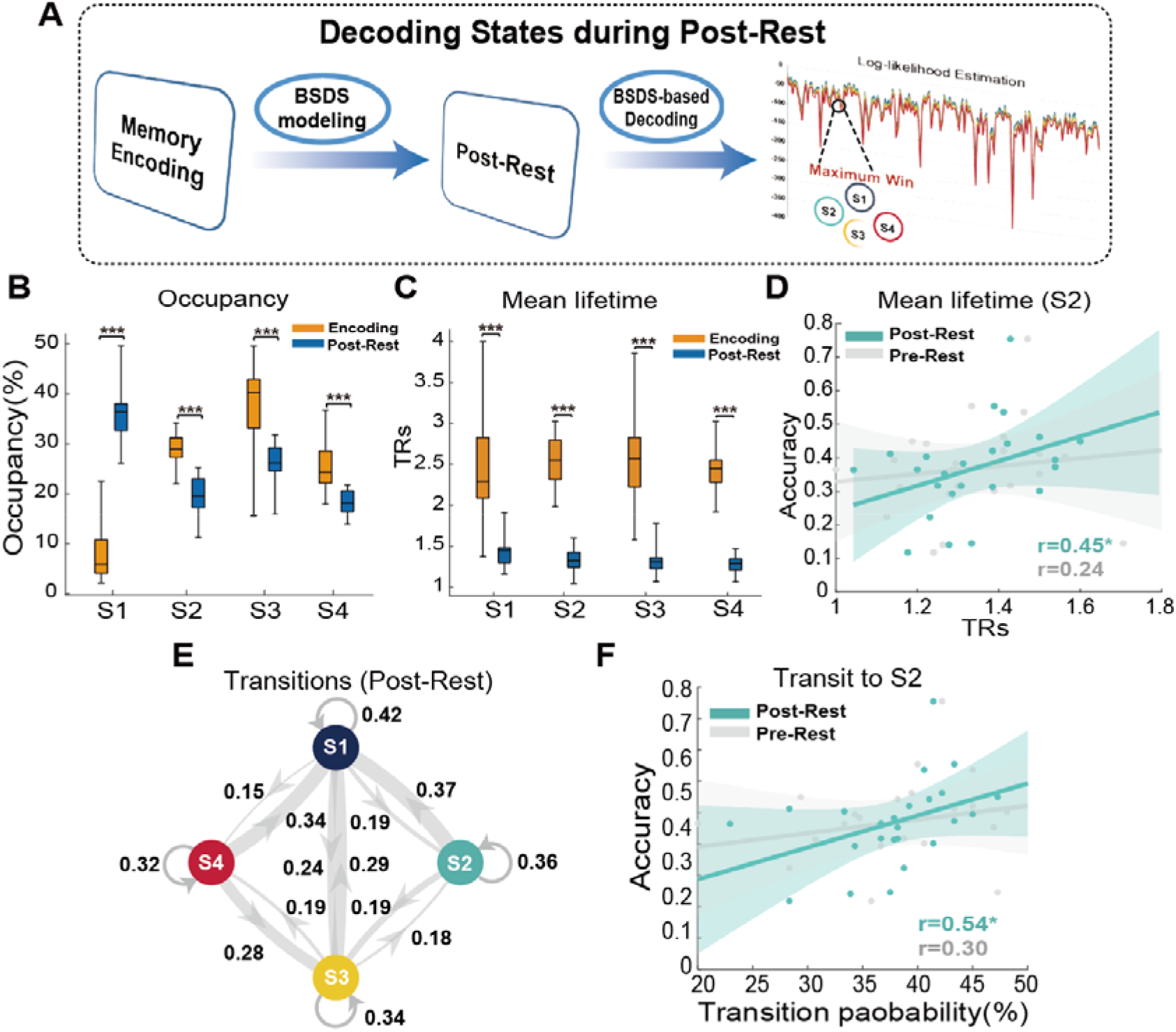
Encoding state reinstatement during post-encoding rest supports memory consolidation. **(A) State decoding framework.** The BSDS model trained on encoding data was applied to post-encoding rest time series via maximum log-likelihood estimation, assigning each TR to the most probable encoding-defined state. **(B) State occupancy shifts from encoding to rest.** The active-encoding state (S3) decreased significantly during rest, while the intermediate state (S1) became the dominant configuration. **(C) Mean lifetime shifts.** All states showed significantly decreased mean lifetimes during post-encoding rest relative to encoding. **(D) Default-mode state reinstatement predicts memory.** Mean lifetime of the default-mode state (S2) during post-encoding rest, but not pre-encoding rest, correlates with memory accuracy, indicating that sustained DMN activity during the post-encoding period supports offline consolidation. **(E) Rest transition dynamics.** Transition probabilities during post-encoding rest show decreased self-transitions and increased inter-state transitions compared to encoding. **(F) Consolidation state stability.** Default-mode state stability predicts memory. Self-transition probability of S2 during post-encoding rest (but not pre-encoding rest) shows the strongest correlation with memory performance, indicating that temporal persistence of this consolidation-related network configuration predicts memory outcomes. Dashed lines indicate 95% confidence intervals; solid lines show best linear fit. Error bars represent SEM. \**p* < 0.05, \*\*\**p* < 0.001.

We first investigated the association of state distribution between encoding and post-encoding rest. By calculating correlations between individual-paired state distributions (**Methods**), we found a significant correlation between the encoding and post-encoding rest phases, when compared with shuffled paired state distributions (r = -0.62, permuted *p* < 0.05), suggesting that such correlation is individual specific (**Supplementary Figure S4A**). Further entropy analyses based on states distribution (Methods) revealed significantly higher entropy during post encoding rest compared with the encoding session (paired t-test, t_(23)_ = 6.32, *p* < 0.001), suggesting a substantial state distribution shift, from a task-prioritized distribution toward a more evenly distributed mode during post-encoding rest (**Supplementary Figure S4B)**.

Next, we compared the state dynamicity shift for each brain state and as indicated in **Fig 4B**, we found that the active-encoding state (S3) which yielded the highest occupancy rate during encoding phase decreased substantially during the post-encoding rest phase (paired t-test: t_(23)_ = -6.02, *p* < 0.001, FDR corrected), and the intermediate state (S1) became the dominant state during post-encoding rest compared with other states (all ts_(23)_ > 6.19, *ps* < 0.001, FDR corrected). In addition, the mean lifetime of all states decreased substantially during post-encoding rest (**Fig. 4C**). Further comparison of state occupancy rate and mean lifetime between pre- and post-rest revealed a significantly higher occupancy rate (t_(23)_ = 2.12, *p* = 0.045, uncorrected) and a marginally higher mean lifetime (t_(23)_ = 1.78, *p* = 0.089, uncorrected) of active-encoding state (S3) during post-encoding rest compared with pre-encoding rest (**Supplementary Figure S5**), suggesting an encoding-induced state dynamic reorganization during post-encoding rest. In terms of transition dynamics, we observed a substantial shift from encoding toward post-encoding rest (**Fig. 4E**), as indicated by a significant Frobenius norm difference between the two transition networks (permuted-*p* < 0.001). But we didn’t observe such shift in transition dynamics between pre- and post-rest (permuted-*p* > 0.28). Together, these results indicate a systematic shift of brain state dynamics from encoding toward a post-encoding reinstatement process.

Finally, we investigated whether the reinstatement of specific encoding states during rest predicted subsequent memory performance. As recent studies have shown that the activity of the default mode network coincide with potential neural replay to further support learning or memory consolidation ^21,38^, we first examined the default-mode state (S2), characterized by both high activation in MPFC and PCC nodes, which showed the strongest reinstatement effects. We found that its mean lifetime during rest correlated robustly with memory accuracy (r = 0.45, *p* = 0.026; **Fig. 4D**), an effect absent for other brain states (all ps > 0.42). Notably, we found such reinstatement effect only occurred during post-encoding rest as occupancy rate in pre-encoding rest did not exhibit similar reinstatement effect (r=0.24, *p* = 0.25), indicating that such reinstatement is specific to post-encoding rest. Furthermore, we investigate whether transitions toward default-mode state are also predictive of memory performance. As expected, we found that the self-transition of default-mode state (S2) during post-encoding rest showed the strongest correlation with memory performance (r = 0.54, *p* < 0.01, FDR corrected; **Fig. 4F**). Again, this memory-predictive effect was absent during pre-encoding rest, confirming specificity to the post-encoding period. Notably, these brain-memory associations were replicated with D-prime as memory performance index (**Supplement Table. S2**).

As an additional control, we trained an independent BSDS model solely on post-encoding rest data to test whether rest-derived states alone could capture the same brain–behavior relationships. This model identified a default-mode state with an amplitude profile significantly similar to the encoding-derived State 2 (Pearson’s r = 0.56, *p* = 0.03), but its dynamics did not predict memory performance (all *ps* > 0.5; **Supplementary Figure S8**), suggesting that encoding-state information is necessary for capturing consolidation-relevant dynamics during post-rest.

To address the potential circularity of deriving ROIs from the subsequent memory contrast, we repeated our analysis pipeline using an independent whole-brain network parcellation ^39^ that was not selected based on memory-related activation (**Methods; Supplementary Figure S9**). This analysis replicated the core findings from both the encoding and post-encoding rest phases, showing that the dynamics of active-encoding states during encoding and the default-mode state during post-encoding rest were associated with memory encoding and reinstatement, further confirming that our findings generalize beyond the original ROI set.

Together, these findings demonstrate that successful episodic memory formation extends beyond initial encoding to include dynamic offline reinstatement processes that strengthen memory representations.

## Discussion

We examined how children’s brains dynamically organize into distinct coordination patterns or “brain states” during memory encoding and how these states are reinstated during rest to support consolidation. Using a novel Bayesian state-space modeling approach, we identified four distinct brain states with unique activation and connectivity signatures during episodic memory encoding scenes. An “active-encoding state” characterized by integrated visual-hippocampal-parietal activity dominated successful encoding. Critically, flexible transitions into the active-encoding state enhanced memory formation. Most importantly, we discovered that encoding states spontaneously reinstated during post-encoding rest, with a “default-mode state” involving mPFC and PCC showing sustained reactivation that robustly predicted memory outcomes. Results reveal that successful memory formation in children depends on dynamic neural flexibility, characterized by the ability to shift into optimal states during learning and to spontaneously reactivate consolidation-related states during rest. These findings advance our understanding of how moment-to-moment brain dynamics bridge encoding and consolidation.

### Dynamic brain states reveal moment-to-moment neural flexibility during the memory encoding phase

We examined how distributed brain regions coordinate their activity over time to support episodic memory formation in children. While extensive evidence demonstrates that memory depends on coordinated interactions between MTL and distributed cortical networks ^10,13,40–43^ – conventional static connectivity approaches cannot capture the dynamic network reconfigurations underlying successful encoding. Recent work has demonstrated that rapid reconfiguration of large-scale brain networks supports cognitive functions including working memory and episodic encoding^26,44,45^. Here we used dynamic state-space hidden Markov models ^31^, and identified four distinct brain states with unique activation and connectivity signatures that emerge during memory encoding. Moreover, state dynamics and inter-state transitions predict memory outcomes, suggesting system-level coordination across distinct systems.

Our core findings that a specific “active-encoding state” characterized by enhanced activity in visual, hippocampal and frontoparietal networks dominates successful encoding and strongly predicted individual memory performance provides direct evidence that optimal network configurations facilitate memory formation. This aligns with extensive evidence showing that visual cortical and hippocampal activity during scene encoding predicts better memory recall ^46,47^, and that lateral frontoparietal coordination guides selective attention to information that the hippocampus binds into relational memory traces ^12,13,48,49^ ^50,51^. Our findings thus provide dynamic network-level evidence for established component-process and attention-to-memory models ^10,18,52^. Critically, the “active-encoding state” state exhibited high global efficiency and clustering coefficient, indicating an optimal balance between network integration and specialized processing. This architecture facilitates efficient binding of visual features into coherent episodic representations, consistent with models proposing that fluctuations between integration and segregation, particularly in frontoparietal-hippocampal systems, support transitions between perceptual processing and relational binding ^31,40,53^.

Notably, we identified two additional brain states with distinct functional profiles. An inactive state characterized by uniformly low amplitude across all regions showed strong negative correlations with memory performance, revealing that successful encoding depends not only on achieving optimal configurations but also on avoiding maladaptive states that impair memory formation. Notably, we detected a default-mode state involving selective activation of mPFC and PCC, core nodes of the DMN known for their role in internal mentation, self-referential processing, and memory integration^20,21^. The emergence of this state during encoding is consistent with recent theoretical frameworks proposing that DMN regions undergo flexible reconfiguration, transiently integrating with task-positive networks to support contextual binding and encoding processes^21,54^. This challenges the traditional view of the DMN as solely a “task-negative” network and instead demonstrates dynamic, state-dependent involvement in memory formation.

Moreover, the presence of this default-mode state during encoding, coupled with the absence of strict trial-by-trial alignment between states and task events, indicates that encoding dynamics are partly internally organized rather than purely stimulus-driven. This internal organization likely reflects fluctuations in attention, engagement, self-referential processing, and encoding strategies that critically determine memory outcomes ^31,55^ ^20,56,57^. Together, these findings demonstrate that dynamic brain states reflect the full spectrum of cognitive processes determining memory outcomes from optimal active-encoding and DMN-mediated configurations to detrimental inactive patterns.

Finally, our analysis of state-to-state transitions revealed another fundamental mechanism: successful encoding depends not only on achieving optimal brain states but also on the flexibility to transition into these states when needed. Transitions toward the active-encoding state from both default-mode and inactive states positively correlated with memory performance, while transitions toward the inactive state were detrimental (**Fig. 3E**). These transition patterns reflect rapid neural reorganization through dynamic assembly of multiple circuits to support learning and flexible information access ^58–60^, consistent with evidence that stable yet flexible hippocampal states maintained by attentional and prefrontal modulation predict successful encoding ^61–63^. This dynamic flexibility extends findings from motor skill learning and working memory showing rapid transitions between network integration and segregation ^45,64,65^, suggesting that memory formation emerges from the brain’s capacity for rapid, adaptive network reorganization rather than sustained activation of fixed circuits.

### Encoding state reinstatement during the post-encoding rest phase reveals offline consolidation mechanisms

Systems consolidation models propose that memory traces are spontaneously reactivated during rest to facilitate long-term retention ^11^. Our state space modeling revealed two key findings during the post-encoding rest. First, all four encoding states spontaneously emerged during rest, with the active-encoding state showing significant increases during this offline period. This indicates that encoding-related network configurations are not simply task-driven responses but represent stable coordination patterns that persist beyond the task context. Second, the “default-mode” state, characterized by enhanced mPFC and PCC activity, demonstrated the strongest reinstatement effects: its mean lifetime during post-encoding rest, a measure of how long this network configuration was sustained each time it was entered, robustly predicted memory performance. That sustained maintenance, rather than mere frequency of occurrence, predicted memory outcomes suggest that the consolidation benefit depends on the temporal stability of DMN-mediated processing during rest. These findings provide direct evidence for systems consolidation through state-specific reinstatement, extending beyond previous studies that focused on regional reactivation patterns ^66,67^. The dissociation between encoding-dominant and rest-dominant states further reveals complementary mechanisms: the active-encoding state supports initial memory formation through sensory-hippocampal integration, while the default-mode state supports offline consolidation through sustained mPFC-PCC activity. This temporal specialization suggests that different network configurations are optimized for different phases of memory, requiring coordinated transitions between states rather than simple persistence of encoding patterns.

This principle aligns is supported by converging evidence. Sustained hippocampal-neocortical coupling increases during post-encoding rest ^68^ and hippocampal replay events co-occur with strong DMN hub activation ^38^, suggesting that successful consolidation relies on coordinated interactions between memory encoding systems and DMN-mediated integration networks ^69^. Our findings extend this framework by demonstrating state-level dynamics: the default-mode state that appears transiently during encoding, where it likely supports contextual integration, becomes substantially more prominent and sustained during offline rest, where it orchestrates memory stabilization and consolidation. This dual, temporally specialized role of DMN regions reconciles seemingly contradictory findings about DMN involvement in memory. Rather than being purely “task-negative,” DMN regions undergo flexible, state-dependent reconfiguration (Menon, 2023): transiently integrating with task-positive encoding networks when contextual binding is needed, then dominating during offline periods to orchestrate sustained consolidation through mPFC-PCC coordination. Indeed, our observed reinstatement and sustained activation of the default-mode state during post-encoding rest provides direct empirical support for this framework, demonstrating that DMN-mediated network dynamics are fundamental for memory stabilization, integration, and adaptive updating during offline consolidation ^21,29,30^.

## Limitations

The current study has several limitations. First, although we aimed to examine whether established state-level consolidation mechanisms operate during childhood, we lacked a direct comparison with adult participants. A direct comparison using the same framework is needed to characterize developmental trajectories of brain state dynamics underlying episodic memory. Second, the relatively small sample size (N = 24) and narrow age range (8–13 years) limited our ability to detect age-related changes in brain state dynamics within our developmental sample. Third, the similarity between old and new test images was not parametrically manipulated or quantified, which may have introduced variability in memory discrimination assessment.

## Conclusions

Our study advances understanding of episodic memory by revealing how dynamic brain state sequences bridge online encoding with offline consolidation. We demonstrate that successful memory formation requires: (1) achieving optimal network configurations during encoding, (2) flexible transitions between states based on cognitive demands, and (3) systematic reinstatement of consolidation-related states during rest. These findings establish dynamic flexibility of brain states as a core principle of memory formation and provide a new framework for understanding how moment-to-moment brain state dynamics support learning and memory in the developing brain. This work opens new avenues for understanding how dynamic neural processes support learning and memory in the developing brain.

## Methods

### Participants

A total of 29 healthy children (16 girls and 13 boys) participated in this study. They were recruited from the San Franciso Bay Area. All participants were right-handed, had no history of neurological or psychiatric diseases, and were not currently taking any medications. To minimize age-related variability, children were selected from an age range of 8 to 13 (mean age: 11.03 ± 1.52). All participants had intelligence quotients (IQ) above 95 and below 145, as measured by the Wechsler Abbreviated Scales of Intelligence (WASI). Verbal, performance and full-scale IQ scores were normalized according to each participant’s age. The study protocol was approved by the Stanford University Institutional Review Board. Written informed consent was obtained from each participant as well as the child’s legal guardian prior to their participation. Participants with root mean squared head motion (i.e., RMS) exceeding one voxel’s width during MR scanning were excluded from further analyses (5 participants), resulting in 24 child participants in the final study analysis.

### General experimental procedures

Participants performed a scene-based episodic memory task ^70,71^. Scenes were chosen because they are known to recruit distributed networks visual cortex, MTL, and frontoparietal regions, providing the multi-region coverage needed for BSDS state modeling. At the start of the experiment each participant underwent a 6-minute pre-encoding resting state scan, and was instructed to close their eyes, let their minds wander. Following this scan each participant performed one episodic memory encoding task during fMRI scanning, where participants were instructed to imagine themselves in each scene and were asked to press a button indicating whether the picture was of a building or a natural scene. Participants then underwent a post-encoding resting scan before the memory test. Outside of the MRI scanner participants performed an episodic memory retrieval task (**Fig. 1A-B**). Details of the two memory tasks are described below. Participants underwent an event-related fMRI while performing the memory encoding task. The event-related fMRI experiment was designed to examine changes in trial-by-trial neural pattern stability with development in terms of the “overlapping waves” model ^72^. The task consisted of 100 trials in total. Each trial consisted of an image presented at the center of screen for 3 seconds, followed by a fixation period jittering from 2.0 to 6.0 seconds. Half of the trials consisted of images of buildings and half of images of natural scenes. The order of the 100 building and natural scene trials was pseudo-randomized across participants with the standard addition and control additional problems always interleaved by a low-level fixation period. Participants performed two experimental runs of the memory encoding task (we combined these two encoding tasks into one and assigned it as ‘the encoding phase’). The length of each experimental run was 6 mins and 30 secs. The other settings were identical with the block design fMRI task.

### Retrieval task

In the memory retrieval task, participants viewed 200 images of buildings and natural scenes, 100 of which they had studied during the memory encoding task and 100 of which they had not. Each trial consisted of an image presented at the center of screen for 3 seconds during which participants indicated via a “Yes” or “No” button press whether or not they had previously studied the picture during the encoding task. The following screen asked participants to indicate, again via a button press, how sure they were of their choice, using the following scale: “Very,”, “Sure,” “A little,” “Unsure.” A fixation period jittered from 2.0 to 6.0 seconds followed. As in the encoding task, half of the trials consisted of images of buildings and half of images of natural scenes. Participants performed four experimental runs of the memory retrieval task. The length of each experimental run was 6 mins and 30 secs. For each participant, we computed the rates of “hits,” calculated as the proportion of studied images that were correctly identified as being seen during the encoding task; “misses,” proportion of studied images that were incorrectly identified as being novel or unseen during the encoding task; “false alarms,” proportion of unstudied images incorrectly identified as studied; and “correct rejections,” proportion of unstudied images correctly identified as novel. From these rates, we also computed hit and miss as a function of confidence (**Supplement Fig. 1**) and participant’s overall memory accuracy as the proportion of correct responses (hits and correct rejections) minus incorrect responses (misses and false alarms). We also implemented D-prime as memory performance to validate the robustness of our observed correlations (**Supplement Table. S2**).

### fMRI data acquisition

Whole brain functional images were acquired from a 3T GE Signa scanner (General Electric, Milwaukee, WI) using a custom-built head coil with a T2*-sensitive gradient echo spiral in-out pulse sequence based on blood oxygenation level-dependent (BOLD) contrast (12). Twenty-nine axial slices (4.0 mm thickness, 0.5 mm skip) parallel to the AC–PC line and covering the whole brain were imaged with the following parameters: volume repetition time (TR) 2.0 sec, echo time (TE) 25 ms, 80° flip angle, matrix size 64 × 64, field of view 200 x 200 mm, and an in-plane spatial resolution of 3.125 mm. To reduce blurring and signal loss arising from field inhomogeneity, an automated high-order shimming method based on spiral acquisitions was used before acquiring functional images. A linear shim correction was applied separately for each slice during reconstruction using a magnetic field map acquired automatically by the pulse sequence at the beginning of the scan.

### fMRI data preprocessing and time series extraction

Images were preprocessed using Statistical Parametric Mapping (SPM8, http://www.fil.ion.ucl.ac.uk/spm). The first eight volumes were discarded for stabilization of the MR signal. Remaining functional images were realigned to correct for rigid-body motion. Subsequently, images were slice-timing corrected, normalized into a standard stereotactic space, and resampled into 2 mm isotropic voxels. Finally, images were spatially smoothed by convolving an isotropic 3D-Gaussian kernel (6-mm full width at half maximum). We selected candidate regions of interest (ROIs) by contrasting neural activity for encoding trials later remembered with later forgotten. First-level GLMs included head-motion parameters as nuisance regressors, with individual Remembered > Forgotten contrasts entered into a group-level parametric one-sample t-test. Significant clusters were identified using a height threshold of *p* < 0.05 and an extent threshold of *p* < 0.05 with cluster-level FWE correction based on nonstationary suprathreshold cluster-size distributions computed using Monte Carlo simulation. Then, to extract time series for later BSDS model training, we first select candidate region of interests by contrasting neural activity for encoding trials later remembered with later forgotten (**Fig. 1C**). By doing so, we selected 16 ROIs covering MTL systems, visual-related areas and frontal-parietal regions as well as default mode hubs (**Supplementary Table. S1**). Each ROI was a 5-mm radius sphere centered at the corresponding peak voxel of each node. For encoding and post-encoding rest session, time series of the mean was extracted from each ROI and were high-pass filtered at 0.008 Hz after which the white matter and cerebrospinal fluid, as well as 6-head movement signals were further used as regressor to remove out from time series. Finally, the time series was linearly detrended and normalized before conducting model training.

### Bayesian Switching Dynamical Systems (BSDS) to infer brain dynamics during encoding

We implemented an advanced Bayesian-based brain state detection model called BSDS to infer brain state during episodic memory task and decode latent state evolution during subsequent post-encoding rest. This model has been used in several recent studies and has the potential to detect cognitive state and pathology of both animal and human brain signals ^31,32^. A detailed description of the model is provided in our previous study ^31^. BSDS could infer brain state or brain network dynamic configuration given a set of ROIs based on the time series. Its outputs consist of 1) Group-level brain state characterization of the state mean amplitude and covariance matrix across nodes. 2) Individual-level state dynamic metrics include occupancy rate and mean lifetime: while the occupancy rate reflects the proportion of time spent in each state, whereas mean lifetime reflects the average duration of each state visit. 3) Group-level inter-state transitions among brain states, which measure the strength of transitions from one state to another.

### Bayesian Switching Dynamical Systems (BSDS) to decode brain dynamics during pre- and post-encoding rest

A unique feature of BSDS is that it could implement the trained model to decode untrained brain states via log-likelihood estimation. Based on this, we leverage the BSDS model trained on encoding session to investigate the brain state dynamics during pre- and post-encoding rest. We extracted the time series during pre/post encoding rest based on identical ROIs in the encoding phase and the first and last 5 volumes were discarded before model inference for robust state estimation. After the model inference, we implemented the maximum-win principle together with threshold (0.1) for z-scored likelihood to determine the state evolution for pre- and post-encoding rest. Based on the decoded brain state sequence, we computed the occupancy rate and mean lifetime of each brain state during pre- and post-encoding rest. We also quantified transition dynamics between states during rest based on the heuristic transition estimation via observed transition behaviors for each participant during rest session.

### Transition network graph analysis

To depict topographical patterns of each latent state, we conduct graph analysis based on Brain Connectivity Toolbox ^73^. Negative covariance value in each covariance matrix is set as zero before conducting network analyses. We computed global efficiency and clustering coefficient given group-level covariance matrix for each brain state. We conducted a permutation test to determine the statistical significance of differences in network properties between each pair of brain states. Consistent with previous study ^74,75^, we circularly shifted state assignment for each participant and computed the corresponding global efficiency and clustering coefficient respectively 5000 times. Then, we obtained null distribution of difference in global efficiency and clustering coefficient between each paired brain states. Finally, we determined the p-value by dividing the number of times the actual difference exceeded the null distribution of correlations.

### Statistical comparison of transition differences

For transition networks among brain states during encoding and post-encoding rest. We first obtained individual-level estimated transitions between brain states and depicted group-level graph network of transition by averaging each specific transition path on group-level. We excluded average transition values below 10% prior to visualization and the thickness of edges between brain states represents the relative strength of state transitions. To test whether the overall transition structure differed between encoding and post-encoding rest, we applied a within-subject sign-flip permutation test based on the Frobenius norm of the group-averaged difference matrices. For each participant, we calculated the difference between transition matrices. The test statistic was defined as the Frobenius norm of the group-averaged difference matrix, representing the overall magnitude of transition change. Then we generated null distribution via randomly flipped each participant’s difference matrix corresponding to a random exchange of condition labels before calculating Frobenius norm for the group-averaged difference matrices. The *p*-value was determined as the proportion of permuted statistics greater than or equal to the observed value (permutation times: 5000).

### Decoding Dominance Value

For determining the decoding performance of state during rest phase, we quantified the strength of evidence of the chosen state over other states. We first defined the Decoding Dominance Value (DDV) for each TR

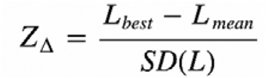

where *L*_best_ and *L*_mean_ denote the highest and mean log-likelihood values across all candidate states at each TR, respectively. And *SD*(*L*) is the standard deviation of the log-likelihood values across all four candidate states at each TR during rest phase. We further construct null distributions by shuffling the log-likelihood values of all four states at each TR for each subject, respectively (permutation times: 1000). We then compared the average DDV across subjects with the null distribution, with the permutation p-value calculated as the proportion of permuted DDV values exceeding the observed average DDV.

### State-memory correlation analyses

To test the association between brain state dynamics and corresponding individual memory performance, we conduct spearman correlation analyses for each brain state metric (occupancy rate/ mean lifetime/ state-to-state transitions) and memory accuracy both during the encoding phase and pre- and the post-encoding rest phase.

### Entropy analysis

To quantify systematic changes in brain state dynamics from encoding to post-encoding rest, we computed the entropy value based on distribution of state occupancy rates as a representative metric. According to information theory^76,77^, entropy measures the uncertainty of a stochastic system—here, a more evenly distribution of states corresponds to higher entropy, whereas a more skewed distribution indicates lower entropy. In the present study, we calculated the entropy based on the probability of each state’s occupancy rate by the following formula:

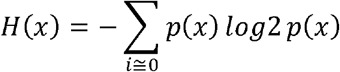

Where p(x) is the probability of each brain state’s occupancy.

### Individual-paired state distribution correlation

To investigate the subject-specific association in states distribution between encoding and post-encoding rest, we conducted a group-level correlation analysis. We first concatenated the state distributions (represented by their occupancy rate) across all subjects for both the encoding phase and their pairing in the post-encoding rest phase. We then computed the correlation between the two phases. To avoid collinearity (because OR’s sum is equal to 1), we included only the first 3 brain states in the analysis. Statistical significance was assessed via a permutation test. To be specific, we randomly shuffle the pairing between encoding phase and post-encoding rest and compute the state distribution correlations 5000 times, thereby generating a null hypothesis distribution. P-value was determined by dividing the number of times the actual correlation is lower than the correlations in the null distribution.

### Robustness analysis

We have conducted 3 distinct parallel BSDS analyses to test the robustness of our result. First, we tested whether BSDS could effectively probed state dynamicity based on event-related design. We shuffle the time series vector of BOLD signal at each TR during encoding phase before parallel BSDS training, as such shuffling could preserve the co-activation pattern across ROIs while disrupting event-related temporal structure (**Supplementary Figure. S7**). Second, we tested whether BSDS model only trained on post-rest data could probed similar brain-behavior correlations. We trained an independent BSDS model based on untrimmed post-rest time series (TRs = 180). The ROIs and hyperparameters used for BSDS training remain unchanged (**Supplementary Figure. S8**). Third, we validate whether the state-behavior correlation could be replicated without resorting to selection that is only based on the subsequent memory contrast. We selected a 14 whole-brain networks template that has been widely used in previous studies ^39^. We exclude 4 of 14 networks before the BSDS modeling: Auditory, Language and motor networks as irrelevant component and primary visual network as redundant. The left 10 network templates are used for extracting time series during encoding and BSDS modeling with the same hyper-parameters setting with the original result (**Supplementary Figure. S9**). Notably, The Central Executive Network (CEN) in this template is referred to as the Frontoparietal Network (FPN) in the subsequent analyses. Finally, permutation test is implemented in determining the significance of correlations between brain dynamics and corresponding memory performance in whole-brain analyses (permutation times: 1000).

## Data availability

All data and codes used to generate the results are available at https://github.com/YimengZeng/Children_Memory_Dynamics.

## Acknowledgements

This research was supported by a National Institutes of Health Career Development Award to SQ (K99MH105601), the National Institutes of Health to V.M. (MH084164, MH126518, HD094623), and the National Natural Science Foundation of China to SQ (32130045) and the Humanities and Social Sciences Fund of the Ministry of Education of China to Y.Z. (25YJC190003).

## Supplementary Information

### I. Supplementary Tables

**Table S1.**
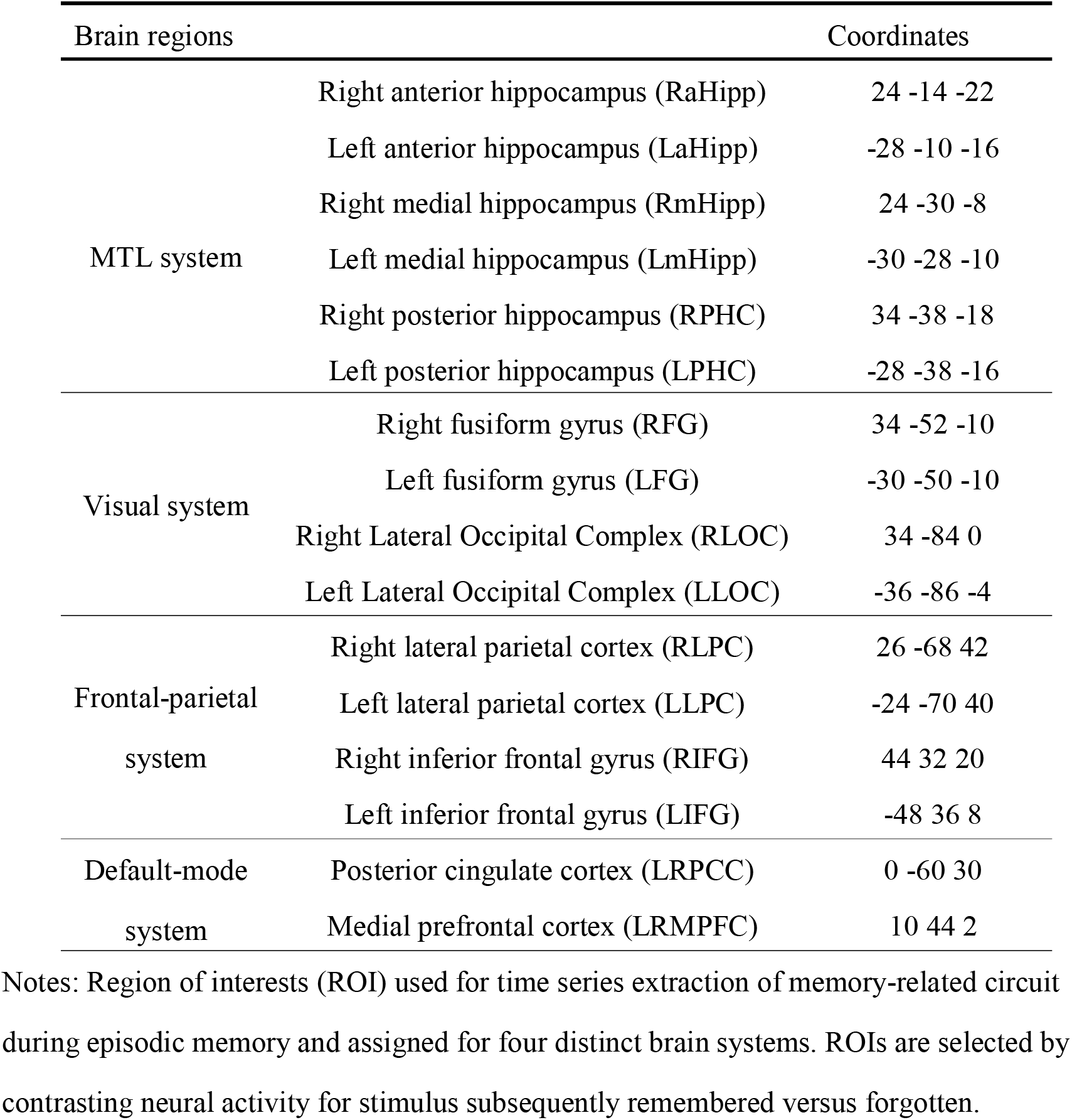
Episodic-memory related regions of interest (ROIs) for modeling.

**Table S2.**
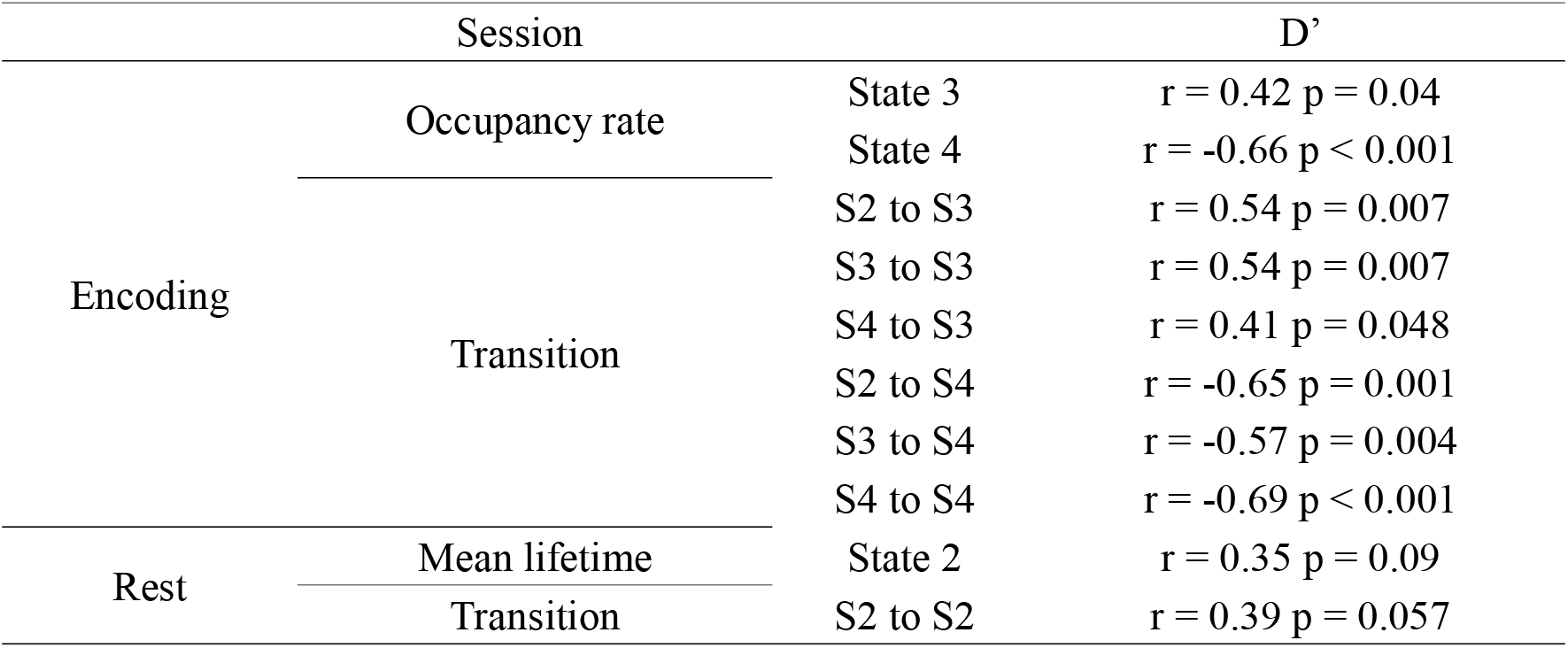
Correlation results using d′ as the measure of memory performance.

### II. Supplementary Figures

**Supplementary Figure S1.**
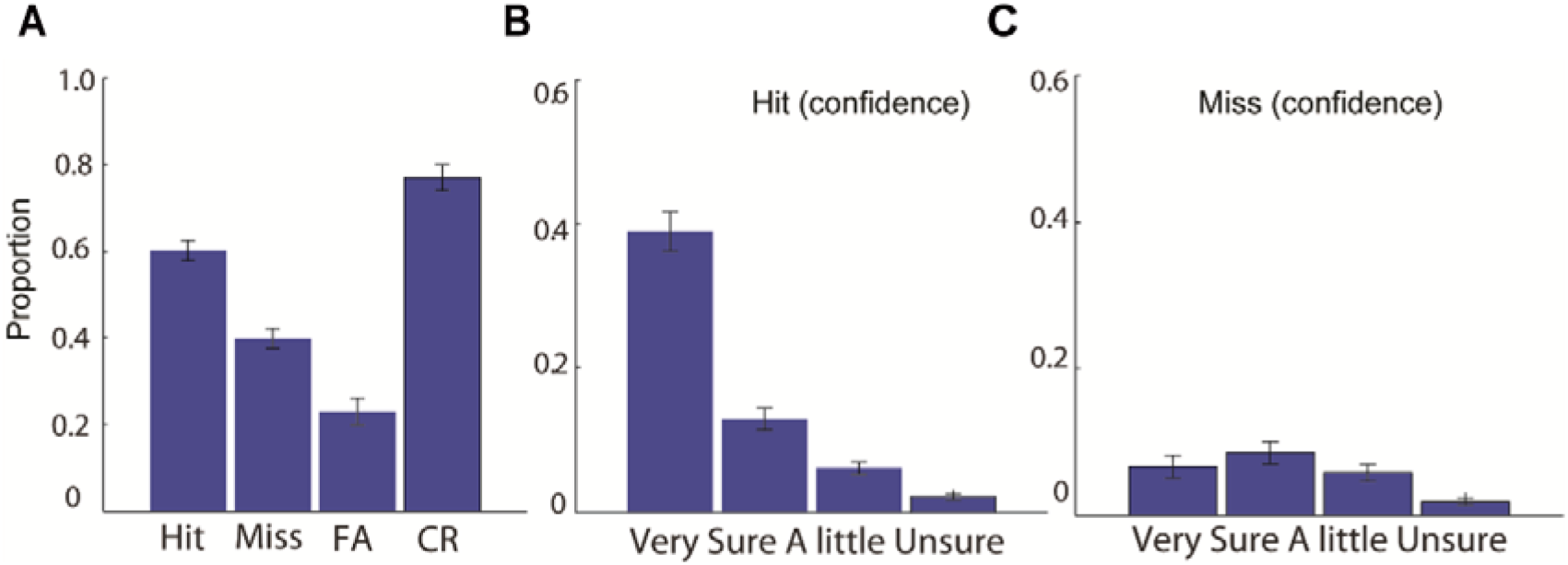
Memory performance validation. **(A-B) Recognition memory results.** Bar graphs showing hit and miss rates as a function of confidence levels. Children demonstrated robust memory performance with hit rates significantly above chance and higher confidence for remembered versus forgotten items. One-way ANOVA revealed significant main effect of confidence ratings, confirming that confidence ratings reliably index memory strength. Error bars represent SEM.

**Supplementary Figure S2.**
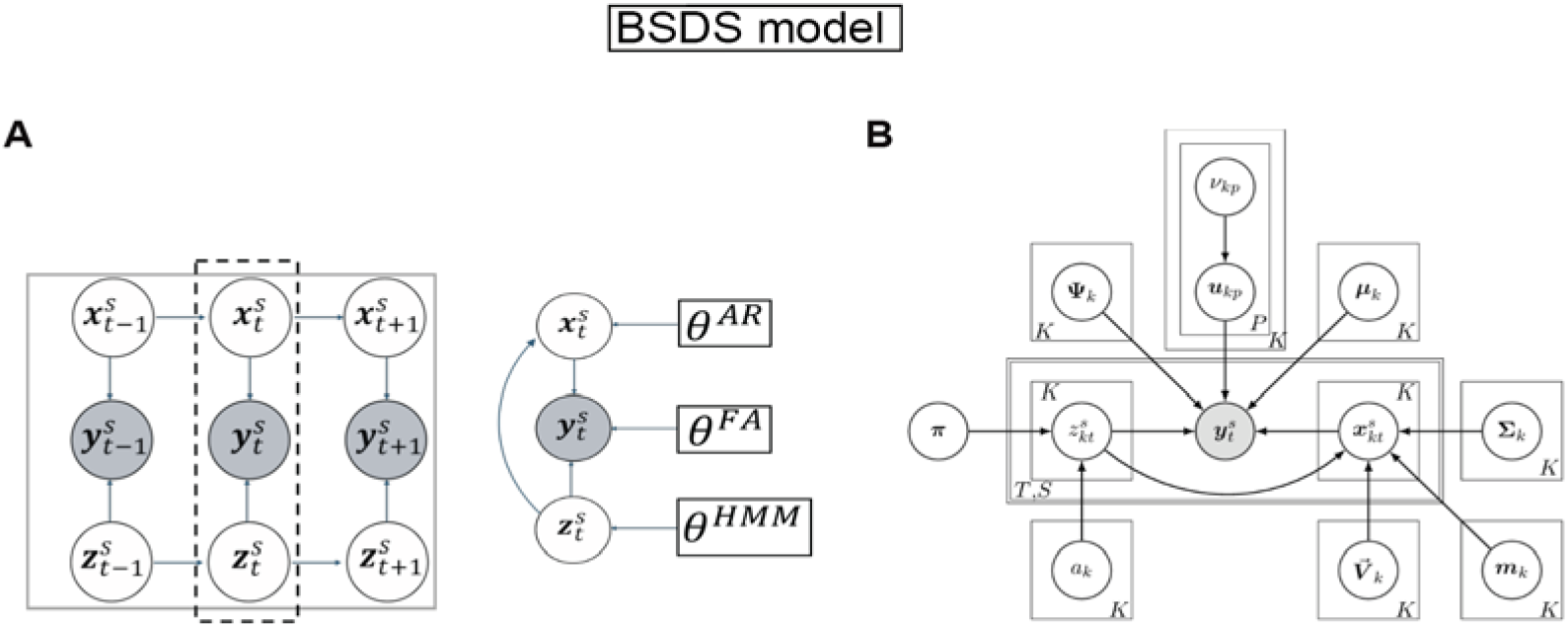
The Bayesian switching dynamical systems (BSDS) model. **(A)** Graphical representation of the BSDS model. ***Y^s^_t_*** is the observed vector of ROI time series at time ***t*** for subject ***s***, ***Z^s^_t_*** denotes the latent states that are interdependent through a first-order Markov chain constrained by a hidden Markov model (HMM) parameterized by ***θ^HMM^***. In any given state, observations ***Y^s^_t_*** are assumed to be generated from a factor analysis (FA) model where ***θ^FA^*** indicates FA model parameters and ***X^s^_t_*** are the latent space variables (factor source variables) of the model. Latent variables are further assumed to be generated using an autoregressive (AR) model, where ***θ^AR^*** indicates the AR model parameters. (**B**) Directed acyclic graph representing the Bayesian switching dynamic systems model. ***Y^s^_t_*** is the observed variable, ***Z^s^_t_*** is the latent state variable, ***X^s^_kt_*** and is the latent source factor (latent space variable) associated to the ***k***-th latent state variable at time ***t*** for subject ***s***. Generative model at time ***t*** and for subject ***s*** at a given latent state k, ***Z^s^_kt_***. π and A are HMM parameters indicating the initial state probability distribution and the state transition probability distribution. ***μ_k_*** is the overall bias in the data, ***Ψ_k_*** is the noise covariance matrix at state ***k***, ***u_kp_*** is the p-th column of the factor loading matrix and at the ***k***-th state, and ***v_kp_***is the prior on the variance of the ***u_kp_***, known as ARD prior which controls the dimensionality of the latent subspace. *ν⃗_k_* indicates the autoregressive coefficients at the ***k***-th state.***Σ_k_*** and **m_k_**are the covariance and the mean in the latent space.

**Supplementary Figure S3.**
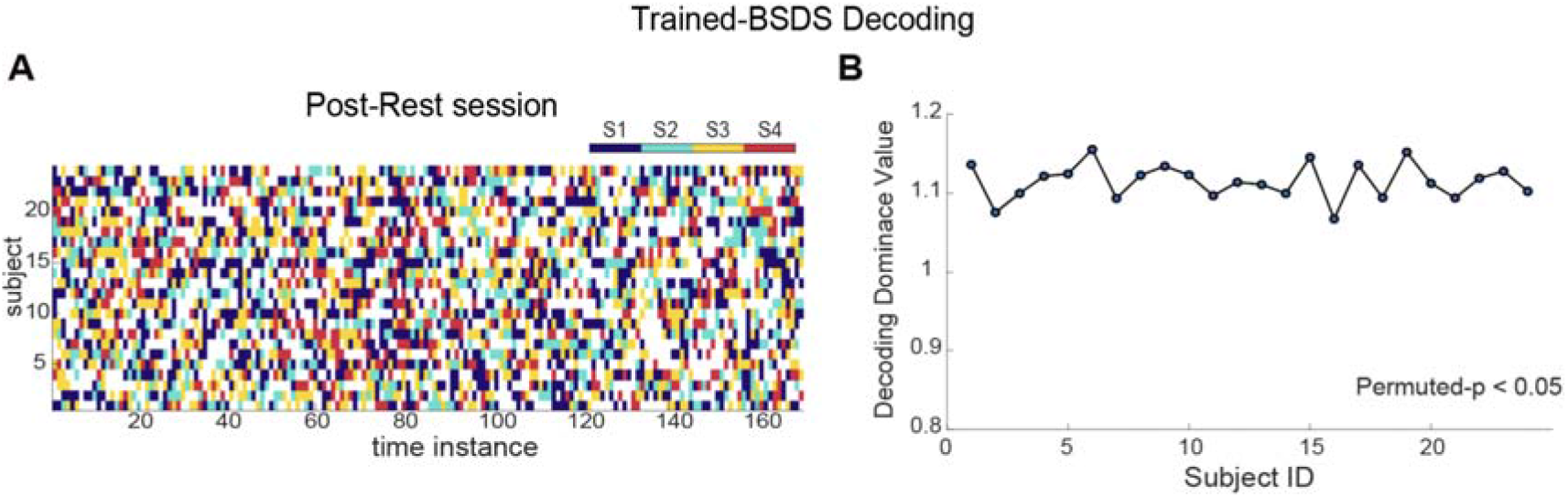
Temporal evolution of brain states. (**A**) State evolution during post-encoding rest decoded using the encoding-trained BSDS model. (**B**) Average Decoding Dominance Value (DDV) for each participant during post-encoding rest, quantifying the strength of evidence favoring the assigned state over alternatives. The observed group mean DDV was significantly higher than the null distribution (permutation *p* < 0.05).

**Supplementary Figure S4.**
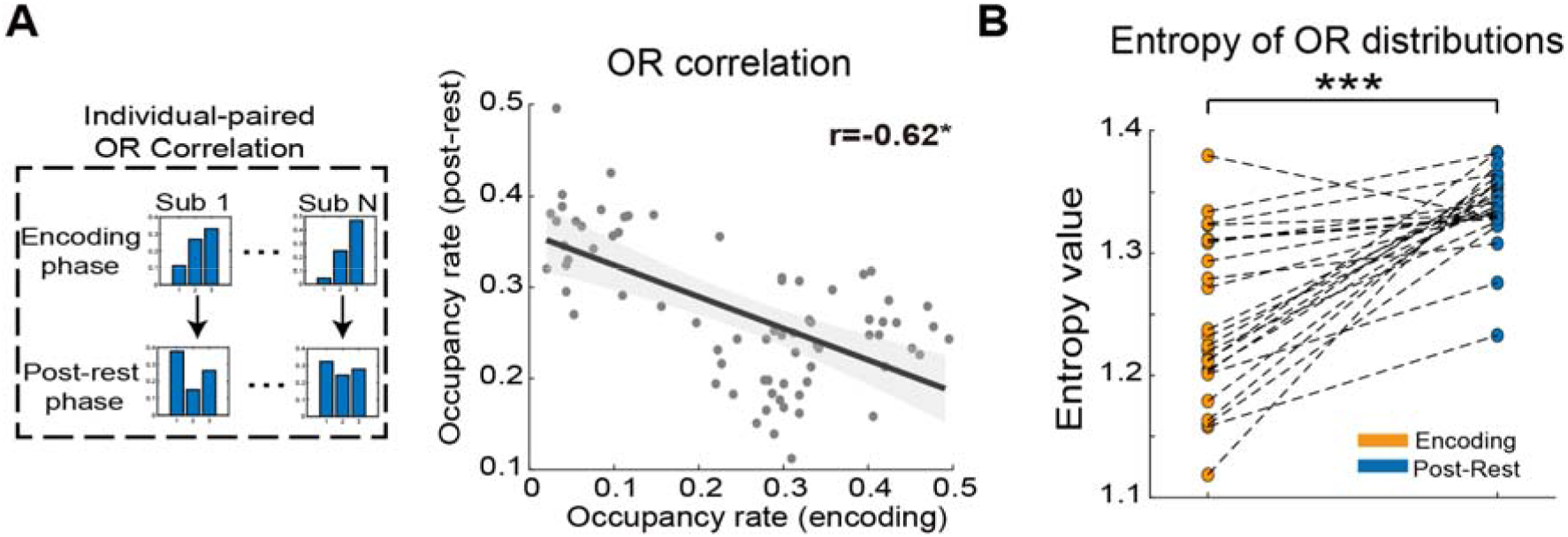
Individual-specific association of brain state distributions between encoding and post-encoding rest. **(A)** Correlation between individual-paired state occupancy rate (OR) distributions across encoding and post-encoding rest. The observed correlation was significantly stronger than expected under shuffled pairings (permutation *p* < 0.05). Dashed lines indicate 95% confidence intervals; solid lines show linear fit. **(B)** Shannon entropy of state occupancy distributions during encoding and post-encoding rest. Higher entropy during post-rest indicates a shift from a task-dominated distribution toward a more uniform state distribution. \**p* < 0.05, \*\*\**p* < 0.001.

**Supplementary Figure S5.**
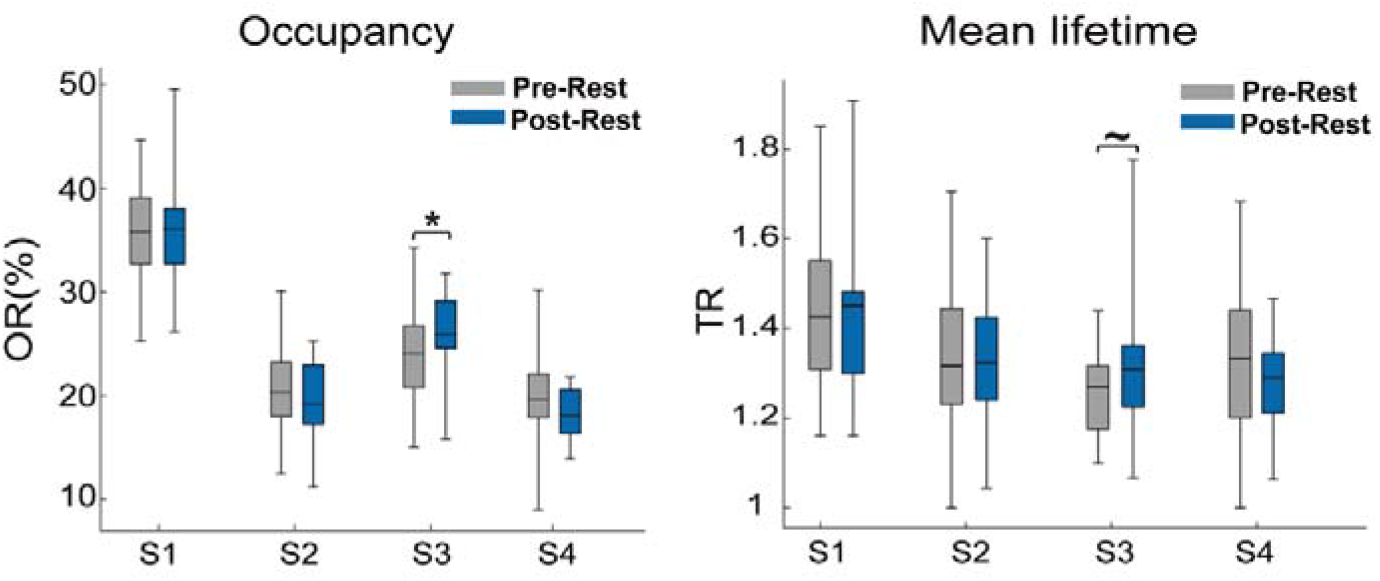
Occupancy rate (OR) and mean lifetime difference between pre- and post-encoding rest. (Left) State occupancy comparison. Box plot comparing state occupancy rate between pre- and post-rest phases. **(Right) Mean lifetime comparison.** Box plot comparing mean lifetime between pre- and post-rest phases. ∼ *p* <0.10, * *p* < 0.05.

**Supplementary Figure S6.**
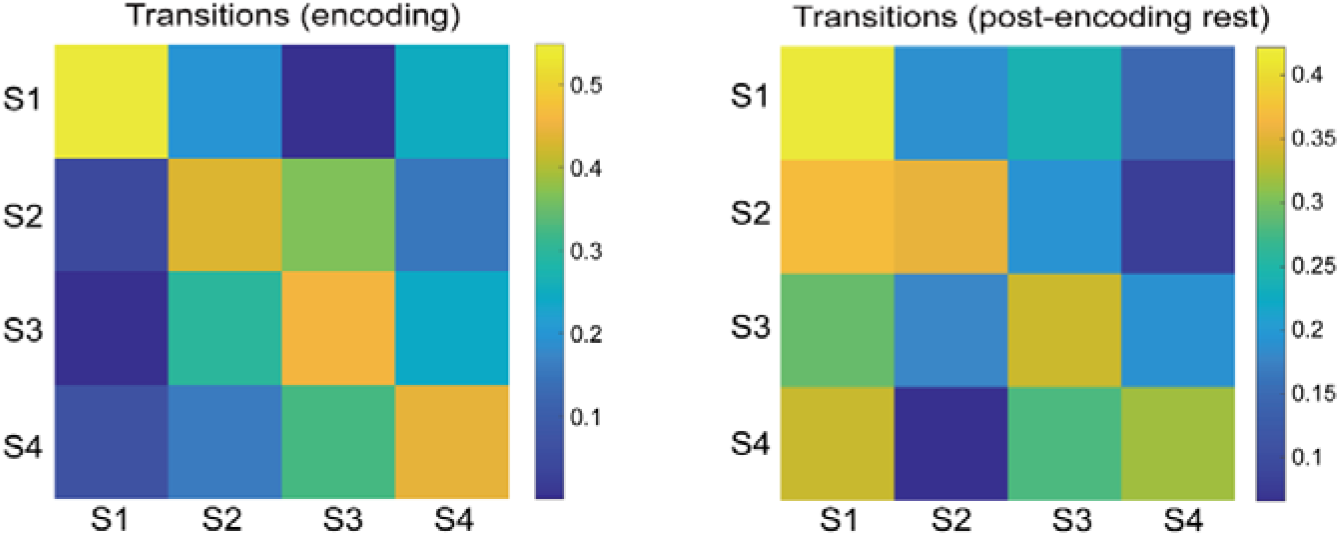
State transition matrices. (Left) Transition matrix for encoding phase. Transition probability matrix shows inter-state dynamics during memory encoding, derived from individual-level BSDS state sequences. Prominent transitions occur between active-encoding (S3), high-order control (S2), and inactive (S4) states. **(Right) Transition matrix for post-encoding rest.** Transition dynamics during post-encoding rest, revealing altered patterns with increased inter-state transitions and decreased self-transitions compared to encoding, reflecting the shift to more variable rest-state dynamics.

**Supplementary Figure S7.**
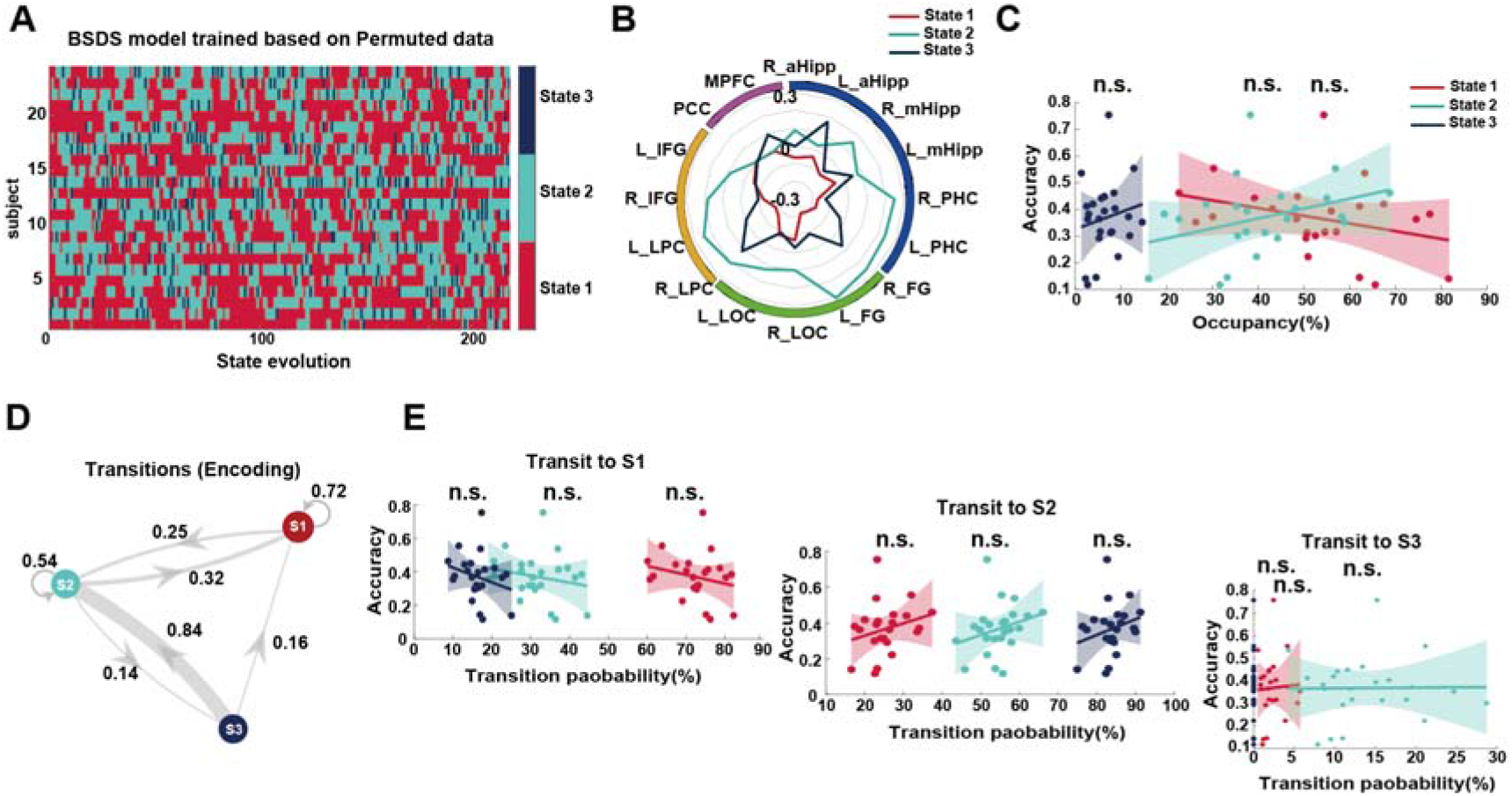
Control analysis: BSDS applied to temporally permuted encoding time series. **(A)** Brain state evolution from BSDS applied to time-shuffled encoding data (first-half TRs shown). Temporal permutation preserved spatial co-activation patterns but disrupted event-related temporal structure. **(B)** Mean amplitude profiles of the three states identified in the permuted data. **(C)** No significant correlations between state occupancy and memory accuracy (all *ps* > 0.24, FDR-corrected). **(D)** Transition network for the permuted data. **(E)** No significant correlations between transition probabilities and memory accuracy (all *ps* > 0.91, FDR-corrected), confirming that the original brain–behavior relationships depend on intact temporal structure. n.s., not significant.

**Supplementary Figure S8.**
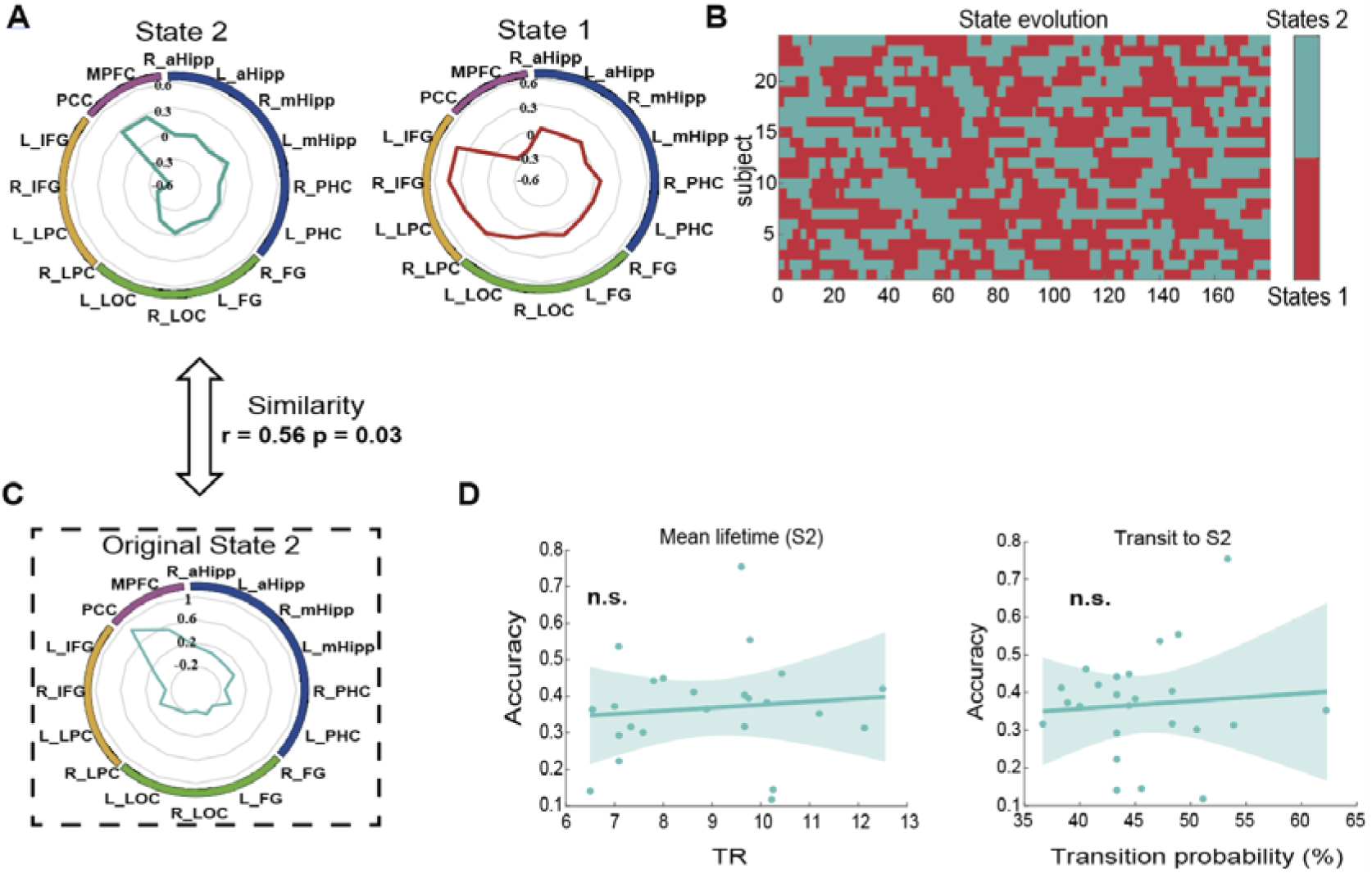
Independent BSDS model trained on post-encoding rest data. **(A)** Two brain states identified by BSDS when trained solely on post-encoding rest time series. **(B)** State evolution. Color-coded temporal sequence of the two states across all participants during post-encoding rest. **(C)** Spatial similarity between the rest-derived State 2 and the encoding-derived default-mode state (S2), confirming significant correspondence in amplitude profiles (r = 0.56, *p* = 0.03). **(D)** Brain–behavior correlations. Neither mean lifetime nor self-transition probability of the rest-derived State 2 predicted memory accuracy (all *ps* > 0.5), suggesting that encoding-state context is necessary for capturing consolidation-relevant dynamics. n.s., not significant.

**Supplementary Figure S9.**
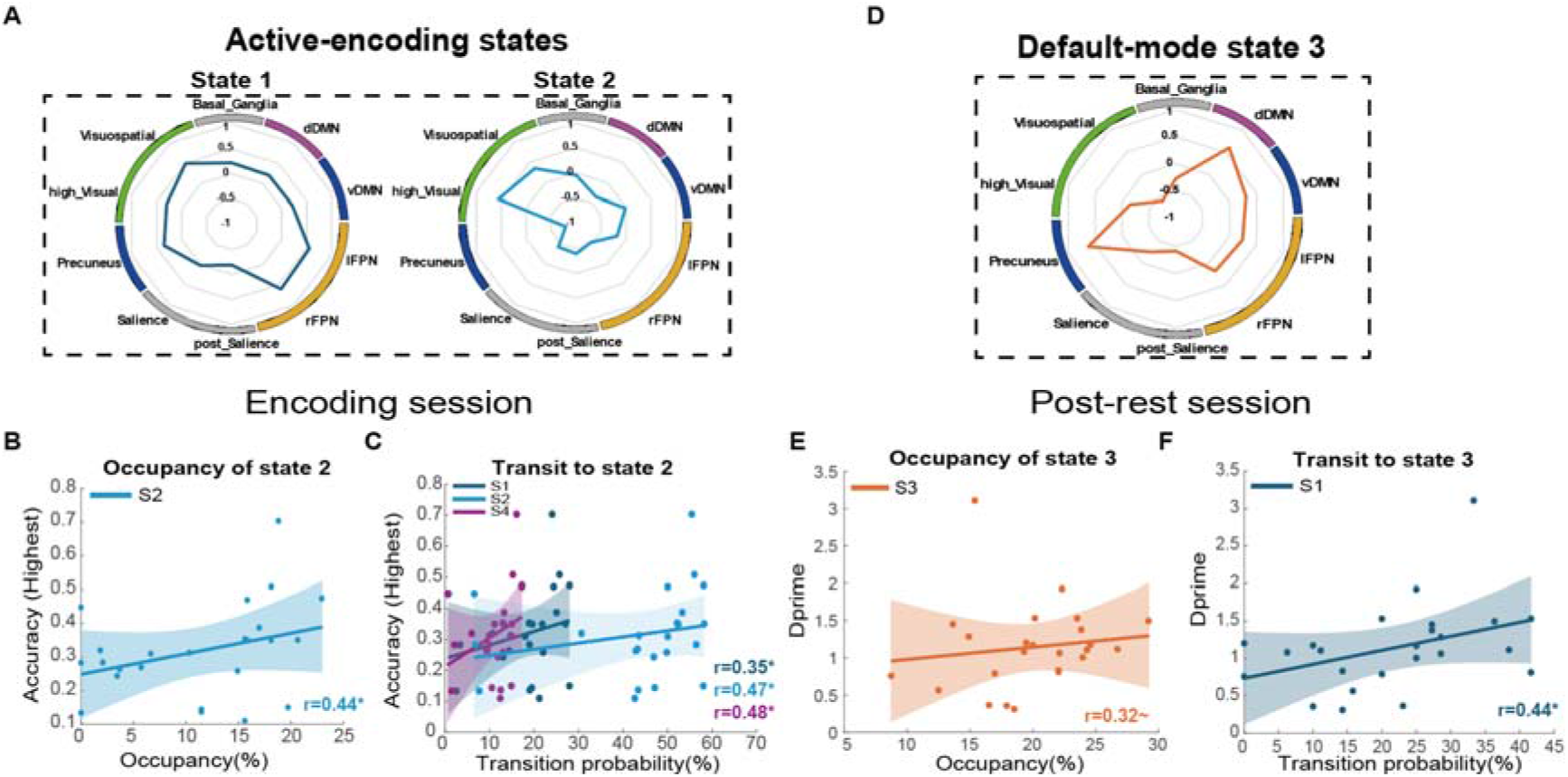
Whole-brain validation using independent network parcellation. **(A)** Mean amplitude of BSDS-identified states 1 and 2 based on the Shirer et al. whole-brain network parcellation. Both states show high activation in visual and lateral FPN, consistent with the active-encoding state from the original ROI analysis. **(B)** Occupancy of state 2 (active-encoding) correlates positively with memory accuracy. **(C)** Transitions toward state 2 correlate positively with memory accuracy. **(D)** Mean amplitude of state 3, which shows elevated activation in the dorsal default-mode network, resembling the default-mode state from the original analysis. **(E)** Transitions toward state 3 during post-encoding rest correlate with memory performance. Abbreviations: dDMN, dorsal default-mode network; vDMN, ventral default-mode network; lFPN/rFPN, left/right frontoparietal networks. Permutation tests: ∼ *p* < 0.1, * *p* < 0.05.

## References

1 Ghetti, S. & Bunge, S. A. Neural changes underlying the development of episodic memory during middle childhood. Developmental cognitive neuroscience 2, 381–395 (2012).

2 Ghetti, S. & Lee, J. Children’s episodic memory. Wiley Interdisciplinary Reviews: Cognitive Science 2, 365–373, doi:10.1002/wcs.114 (2011).

3 Güler, O. E. & Thomas, K. M. Developmental differences in the neural correlates of relational encoding and recall in children: an event-related fMRI study. Developmental cognitive neuroscience 3, 106–116 (2013).

4 Tulving, E. Episodic Memory: From Mind to Brain. Annual Review of Psychology 53, 1–25, doi:10.1146/annurev.psych.53.100901.135114 (2002).

5 Fernández, G. & Tendolkar, I. The rhinal cortex: ‘gatekeeper’ of the declarative memory system. Trends in Cognitive Sciences 10, 358–362, 10.1016/j.tics.2006.06.003 (2006).

6 Zhu, Y. et al. Emotional learning retroactively promotes memory integration through rapid neural reactivation and reorganization. eLife 11, e60190, doi:10.7554/eLife.60190 (2022).

7 Wimmer, G. E., Liu, Y., Vehar, N., Behrens, T. E. J. & Dolan, R. J. Episodic memory retrieval success is associated with rapid replay of episode content. Nature Neuroscience 23, 1025–1033, doi:10.1038/s41593-020-0649-z (2020).

8 Das, A. & Menon, V. Electrophysiological dynamics of salience, default mode, and frontoparietal networks during episodic memory formation and recall revealed through multi-experiment iEEG replication. Elife 13, RP99018 (2024).

9 Zhuang, L. et al. Rapid neural reorganization during retrieval practice predicts subsequent long-term retention and false memory. Nature Human Behaviour, doi:10.1038/s41562-021-01188-4 (2021).

10 Moscovitch, M., Cabeza, R., Winocur, G. & Nadel, L. Episodic Memory and Beyond: The Hippocampus and Neocortex in Transformation. Annu Rev Psychol 67, 105–134, doi:10.1146/annurev-psych-113011-143733 (2016).

11 Kumaran, D., Hassabis, D. & McClelland, J. L. What Learning Systems do Intelligent Agents Need? Complementary Learning Systems Theory Updated. Trends in Cognitive Sciences 20, 512–534, 10.1016/j.tics.2016.05.004 (2016).

12 Eichenbaum, H. Prefrontal–hippocampal interactions in episodic memory. Nature Reviews Neuroscience 18, 547–558, doi:10.1038/nrn.2017.74 (2017).

13 Ranganath, C. & Ritchey, M. Two cortical systems for memory-guided behaviour. Nature Reviews Neuroscience 13, 713–726, doi:10.1038/nrn3338 (2012).

14 Baldassano, C. et al. Discovering Event Structure in Continuous Narrative Perception and Memory. Neuron 95, 709–721.e705, 10.1016/j.neuron.2017.06.041 (2017).

15 Amer, T. & Davachi, L. Extra-hippocampal contributions to pattern separation. elife 12, e82250 (2023).

16 Uncapher, M. R., Hutchinson, J. B. & Wagner, A. D. Dissociable effects of top-down and bottom-up attention during episodic encoding. Journal of Neuroscience 31, 12613–12628 (2011).

17 Ray, K. L. et al. Dynamic reorganization of the frontal parietal network during cognitive control and episodic memory. *Cognitive, Affective*, & Behavioral Neuroscience 20, 76–90, doi:10.3758/s13415-019-00753-9 (2020).

18 Cabeza, R., Ciaramelli, E., Olson, I. R. & Moscovitch, M. The parietal cortex and episodic memory: an attentional account. Nature Reviews Neuroscience 9, 613–625, doi:10.1038/nrn2459 (2008).

19 He, Y. et al. Development of brain-state dynamics involved in working memory. Cerebral Cortex, doi:10.1093/cercor/bhad022 (2023).

20 Buckner, R. L. & DiNicola, L. M. The brain’s default network: updated anatomy, physiology and evolving insights. Nature Reviews Neuroscience 20, 593–608, doi:10.1038/s41583-019-0212-7 (2019).

21 Menon, V. 20 years of the default mode network: A review and synthesis. Neuron 111, 2469–2487 (2023).

22 Ryali, S. et al. Temporal Dynamics and Developmental Maturation of Salience, Default and Central-Executive Network Interactions Revealed by Variational Bayes Hidden Markov Modeling. PLoS Comput Biol 12, e1005138, doi:10.1371/journal.pcbi.1005138 (2016).

23 Chen, T., Cai, W., Ryali, S., Supekar, K. & Menon, V. Distinct Global Brain Dynamics and Spatiotemporal Organization of the Salience Network. PLOS Biology 14, e1002469, doi:10.1371/journal.pbio.1002469 (2016).

24 Meer, J. N. v. d., Breakspear, M., Chang, L. J., Sonkusare, S. & Cocchi, L. Movie viewing elicits rich and reliable brain state dynamics. Nature Communications 11, 5004, doi:10.1038/s41467-020-18717-w (2020).

25 Stitt, I. et al. Dynamic reconfiguration of cortical functional connectivity across brain states. Scientific Reports 7, 8797, doi:10.1038/s41598-017-08050-6 (2017).

26 Phan, A. T., Xie, W., Chapeton, J. I., Inati, S. K. & Zaghloul, K. A. Dynamic patterns of functional connectivity in the human brain underlie individual memory formation. Nature Communications 15, 8969, doi:10.1038/s41467-024-52744-1 (2024).

27 Zielinski, M. C., Tang, W. & Jadhav, S. P. The role of replay and theta sequences in mediating hippocampal-prefrontal interactions for memory and cognition. Hippocampus 30, 60–72, 10.1002/hipo.22821 (2020).

28 Kitamura, T. et al. Engrams and circuits crucial for systems consolidation of a memory. *Science (New York*, N.Y.) 356, 73–78, doi:10.1126/science.aam6808 (2017).

29 Tompary, A. & Davachi, L. Consolidation Promotes the Emergence of Representational Overlap in the Hippocampus and Medial Prefrontal Cortex. Neuron 96, 228–241.e225, 10.1016/j.neuron.2017.09.005 (2017).

30 Tompary, A. & Davachi, L. Integration of overlapping sequences emerges with consolidation through medial prefrontal cortex neural ensembles and hippocampal–cortical connectivity. Elife 13, e84359 (2024).

31 Taghia, J. et al. Uncovering hidden brain state dynamics that regulate performance and decision-making during cognition. Nature Communications 9, 2505, doi:10.1038/s41467-018-04723-6 (2018).

32 Cai, W. et al. Latent brain state dynamics distinguish behavioral variability, impaired decision-making, and inattention. Molecular Psychiatry, doi:10.1038/s41380-021-01022-3 (2021).

33 de Chastelaine, M. & Rugg, M. D. The relationship between task-related and subsequent memory effects. Human Brain Mapping 35, 3687–3700, doi:10.1002/hbm.22430 (2014).

34 Brewer, J. B., Zhao, Z., Desmond, J. E., Glover, G. H. & Gabrieli, J. D. E. Making Memories: Brain Activity that Predicts How Well Visual Experience Will Be Remembered. Science 281, 1185, doi:10.1126/science.281.5380.1185 (1998).

35 Karlsson, M. P. & Frank, L. M. Awake replay of remote experiences in the hippocampus. Nature Neuroscience 12, 913, doi:10.1038/nn.2344 https://www.nature.com/articles/nn.2344#supplementary-information (2009).

36 Sandrini, M., Cohen, L. G. & Censor, N. Modulating reconsolidation: a link to causal systems-level dynamics of human memories. Trends in Cognitive Sciences 19, 475–482, 10.1016/j.tics.2015.06.002 (2015).

37 Schwabe, L., Nader, K. & Pruessner, J. C. Reconsolidation of Human Memory: Brain Mechanisms and Clinical Relevance. Biological Psychiatry 76, 274–280, 10.1016/j.biopsych.2014.03.008 (2014).

38 Higgins, C. et al. Replay bursts in humans coincide with activation of the default mode and parietal alpha networks. Neuron 109, 882–893. e887 (2021).

39 Shirer, W. R., Ryali, S., Rykhlevskaia, E., Menon, V. & Greicius, M. D. Decoding subject-driven cognitive states with whole-brain connectivity patterns. Cereb Cortex 22, 158–165, doi:10.1093/cercor/bhr099 (2012).

40 Geib, B. R., Stanley, M. L., Dennis, N. A., Woldorff, M. G. & Cabeza, R. From hippocampus to whole-brain: The role of integrative processing in episodic memory retrieval. Human Brain Mapping 38, 2242–2259, 10.1002/hbm.23518 (2017).

41 Cohen, N. J. & Eichenbaum, H. Memory, amnesia, and the hippocampal system. (MIT press, 1993).

42 Squire, L. R. Memory and the hippocampus: a synthesis from findings with rats, monkeys, and humans. Psychological review 99, 195 (1992).

43 Rugg, M. D. & Vilberg, K. L. Brain networks underlying episodic memory retrieval. Current opinion in neurobiology 23, 255–260 (2013).

44 Shine, J. M. & Poldrack, R. A. Principles of dynamic network reconfiguration across diverse brain states. NeuroImage 180, 396–405, 10.1016/j.neuroimage.2017.08.010 (2018).

45 Braun, U. et al. Dynamic reconfiguration of frontal brain networks during executive cognition in humans. Proc Natl Acad Sci U S A 112, 11678–11683, doi:10.1073/pnas.1422487112 (2015).

46 Rosen, M. L. et al. The role of visual association cortex in associative memory formation across development. Journal of cognitive neuroscience 30, 365–380 (2018).

47 Chai, X. J., Tang, L., Gabrieli, J. D. & Ofen, N. From vision to memory: How scene-sensitive regions support episodic memory formation during child development. Developmental Cognitive Neuroscience 65, 101340 (2024).

48 Summerfield, C. et al. Neocortical connectivity during episodic memory formation. PLoS biology 4, e128 (2006).

49 Blumenfeld, R. S. & Ranganath, C. Prefrontal cortex and long-term memory encoding: an integrative review of findings from neuropsychology and neuroimaging. The Neuroscientist 13, 280–291 (2007).

50 Rugg, M. D., Johnson, J. D., Park, H. & Uncapher, M. R. Encoding-retrieval overlap in human episodic memory: a functional neuroimaging perspective. Progress in brain research 169, 339–352 (2008).

51 Tang, L., Shafer, A. T. & Ofen, N. Prefrontal cortex contributions to the development of memory formation. Cerebral Cortex 28, 3295–3308 (2018).

52 Uncapher, M. R. & Wagner, A. D. Posterior parietal cortex and episodic encoding: insights from fMRI subsequent memory effects and dual-attention theory. Neurobiology of learning and memory 91, 139–154 (2009).

53 Xue, G. et al. Greater neural pattern similarity across repetitions is associated with better memory. Science 330, 97–101 (2010).

54 Ferris, C., Scheurich, R., Palmer, C. & Sheldon, S. Hippocampal-cortical networks predict conceptual versus perceptually guided narrative memory. The Journal of Neuroscience, e1936242025, doi:10.1523/jneurosci.1936-24.2025 (2025).

55 Al-Aidroos, N., Said, C. P. & Turk-Browne, N. B. Top-down attention switches coupling between low-level and high-level areas of human visual cortex. Proceedings of the National Academy of Sciences 109, 14675, doi:10.1073/pnas.1202095109 (2012).

56 Mesulam, M. M. Large scale neurocognitive networks and distributed processing for attention, language, and memory. Annals of Neurology: Official Journal of the American Neurological Association and the Child Neurology Society 28, 597–613 (1990).

57 Fornito, A., Harrison, B. J., Zalesky, A. & Simons, J. S. Competitive and cooperative dynamics of large-scale brain functional networks supporting recollection. Proceedings of the National Academy of Sciences 109, 12788–12793 (2012).

58 Ito, J., Nikolaev, A. R. & Leeuwen, C. v. Dynamics of spontaneous transitions between global brain states. Human Brain Mapping 28, 904–913, doi:10.1002/hbm.20316 (2007).

59 Zeng, Y. et al. Cortisol awakening response prompts dynamic reconfiguration of brain networks in emotional and executive functioning. Proceedings of the National Academy of Sciences 121, e2405850121, doi:doi:10.1073/pnas.2405850121 (2024).

60 Wu, M. W., Kourdougli, N. & Portera-Cailliau, C. Network state transitions during cortical development. Nature Reviews Neuroscience 25, 535–552, doi:10.1038/s41583-024-00824-y (2024).

61 Aly, M. & Turk-Browne, N. B. Attention promotes episodic encoding by stabilizing hippocampal representations. Proceedings of the National Academy of Sciences 113, E420–E429 (2016).

62 Kuhl, B. A., Rissman, J. & Wagner, A. D. Multi-voxel patterns of visual category representation during episodic encoding are predictive of subsequent memory. Neuropsychologia 50, 458–469 (2012).

63 Xue, G. The Neural Representations Underlying Human Episodic Memory. Trends in Cognitive Sciences 22, 544–561, 10.1016/j.tics.2018.03.004 (2018).

64 Bassett, D. S. et al. Dynamic reconfiguration of human brain networks during learning. Proc Natl Acad Sci U S A 108, 7641–7646, doi:10.1073/pnas.1018985108 (2011).

65 Shine, J. M., Aburn, M. J., Breakspear, M. & Poldrack, R. A. The modulation of neural gain facilitates a transition between functional segregation and integration in the brain. Elife 7, doi:10.7554/eLife.31130 (2018).

66 Carr, M. F., Jadhav, S. P. & Frank, L. M. Hippocampal replay in the awake state: a potential substrate for memory consolidation and retrieval. Nature Neuroscience 14, 147, doi:10.1038/nn.2732 (2011).

67 Tambini, A. & Davachi, L. Persistence of hippocampal multivoxel patterns into postencoding rest is related to memory. Proceedings of the National Academy of Sciences 110, 19591, doi:10.1073/pnas.1308499110 (2013).

68 Folvik, L. et al. Sustained upregulation of widespread hippocampal–neocortical coupling following memory encoding. Cerebral Cortex 33, 4844–4858, doi:10.1093/cercor/bhac384 (2022).

69 Kaefer, K., Stella, F., McNaughton, B. L. & Battaglia, F. P. Replay, the default mode network and the cascaded memory systems model. Nature Reviews Neuroscience 23, 628–640 (2022).

70 Tambini, A., Rimmele, U., Phelps, E. A. & Davachi, L. Emotional brain states carry over and enhance future memory formation. Nature Neuroscience 20, 271–278, doi:10.1038/nn.4468 (2017).

71 Tambini, A. & Davachi, L. Awake Reactivation of Prior Experiences Consolidates Memories and Biases Cognition. Trends in Cognitive Sciences 23, 876–890, 10.1016/j.tics.2019.07.008 (2019).

72 Siegler, R. S. Emerging minds: The process of change in children’s thinking. (Oxford University Press, 1998).

73 Rubinov, M. & Sporns, O. Complex network measures of brain connectivity: uses and interpretations. Neuroimage 52, 1059–1069, doi:10.1016/j.neuroimage.2009.10.003 (2010).

74 Zeng, Y. et al. Dynamic integration and segregation of amygdala subregional functional circuits linking to physiological arousal. NeuroImage, 118224, 10.1016/j.neuroimage.2021.118224 (2021).

75 Kauppi, J.-P., Jääskeläinen, I., Sams, M. & Tohka, J. Inter-subject correlation of brain hemodynamic responses during watching a movie: localization in space and frequency. Frontiers in Neuroinformatics 4, doi:10.3389/fninf.2010.00005 (2010).

76 Munn, B. R., Müller, E. J., Wainstein, G. & Shine, J. M. The ascending arousal system shapes neural dynamics to mediate awareness of cognitive states. Nature Communications 12, 6016, doi:10.1038/s41467-021-26268-x (2021).

77 Ezaki, T., Watanabe, T., Ohzeki, M. & Masuda, N. Energy landscape analysis of neuroimaging data. *Philosophical Transactions of the Royal Society A: Mathematical*, Physical and Engineering Sciences 375, 20160287, doi:10.1098/rsta.2016.0287 (2017).

